# Structure of the II2-III2-IV2 mitochondrial supercomplex from the parasite *Perkinsus marinus*

**DOI:** 10.1101/2024.05.25.595893

**Authors:** Fēi Wú, Alexander Mühleip, Thomas Gruhl, Lilach Sheiner, Amandine Maréchal, Alexey Amunts

## Abstract

Respiratory complexes have co-evolved into supercomplexes in different clades to sustain energy production at the basis of eukaryotic life. In this study, using cryogenic electron microscopy, we determined the 2.1 Å resolution structure of a 104-subunit II2-III2-IV2 supercomplex from the parasite Perkinsus marinus, related to Apicomplexa, capable of complete electron transport from succinate to molecular oxygen. A feature of the parasite is the association of two copies of complex II via the apicomplexan subunit SDHG that interacts with both complexes III and IV and bridge the supercomplex. In the *c*_1_ state, we identified two protein factors, ISPR1 and ISPR2 bound on the surface of complex III, where Cytochrome *c* docks, acting as negative regulators. The acquisition of 15 specific subunits to complex IV results in its lateral offset, increasing the distance between the Cytochrome *c* electron donor and acceptor sites. The domain homologous to canonical mitochondria-encoded transmembrane subunit COX2 is made of three separate polypeptides encoded in the nucleus, and their correct assembly is a prerequisite for electron transport in the supercomplex. Subunits Cytochrome *b* and COX1 comprise a +2 frameshift introduced during protein synthesis by the mitoribosome. Among 114 modelled endogenous lipids, we detect a direct contribution to the formation of the divergent supercomplex and its functional sites, including assembly of CII and ubiquinone binding. Together, our findings expose the uniqueness of the principal components of bioenergetics in the mitochondria of parasites.

---

Eukaryotes rely on oxidative phosphorylation and the crucial energy conversion provided by encapsulated mitochondria for their aerobic metabolism. This occurs through respiratory chains and the coupled transport of electrons and protons across multi-protein complexes embedded in the inner mitochondrial membrane (IMM), ultimately leading to adenosine triphosphate (ATP) synthesis. While these respiratory complexes were initially characterised individually, it is now evident that many of them can assemble into higher organisational order assemblies termed supercomplexes (SCs). Functionally, SCs can facilitate complete electron transport from nicotinamide adenine dinucleotide (NADH) to molecular oxygen. At least four different stoichiometries and composition types of SCs have been reported *in situ* across various regions of the mammalian mitochondrial membrane ^1^, containing complex I (CI, NADH:ubiquinone oxidoreductase), III_2_ (CIII, ubiquinol:cytochrome *c* oxidoreductase or cytochrome *bc*_1_) and IV (CIV, cytochrome *c* oxidase). To date, no mammalian SC has been identified to contain complex II (CII, succinate:UQ oxidoreductase). In humans, different stoichiometries of SCs have been associated with metabolic state changes ^2,3^ and diseases ^4,5^. In other organisms, SCs can reach a molecular weight of 10 MDa, and recent studies have highlighted their central role in mitochondrial ultrastructure and function ^6–9^.

The respiratory chains of parasites likely exhibit noticeable functional and compositional distinctions, which have not been fully investigated and may be linked to disease. In addition, parasitic microorganisms possess some of the most reduced and divergent mitochondrial DNA ^10,11^, making them intriguing for insights into less conventional modes of life. Particularly, the mitochondria of the infectious Apicomplexa parasites display unique cristae morphology shaped by the ATP synthase cyclic hexamers localised to curved apical membrane regions ^12^. Although these parasites lack CI, proteomic analysis of their CII, CIII and CIV has revealed significant compositional differences, with many detected subunits being previously unknown, suggesting a potentially unique protein organisation ^13,14^. Understanding the principles of electron transport in these complexes is crucial for parasite physiology and potential therapeutic interventions targeting energy production. However, fundamental mechanistic aspects in parasites have remained elusive due to a lack of corresponding structural information, including respiratory complexes organisation. We thus set to determine the atomic structures of noncanonical SCs and identify specific factors underlying their unique characteristics.

Here, we report 2.1-2.4 Å resolution structures of the mitochondrial II2-III_2_-IV2 SC from the parasite *Perkinsus marinus* that infects molluscs and is related to Apicomplexa, using cryogenic-electron microscopy (cryo-EM). We show that, in addition to CIII and CIV, two monomers of functional CII are associated with the SC, resulting in a stoichiometric II2-III_2_-IV2 that has not been observed in any other organism. This SC is capable of electron transport from succinate to oxygen, and represents a new type of functional respirasome. The data enabled visualization of native lipids with key roles in SC formation and activity. Finally, we identified an associated heterodimer factor as a potential regulator of CIII locking the Rieske iron-sulfur protein (RIP1) head in the *c*_1_-state, which we named Iron-Sulfur Protein Related (ISPR1,2).

## Results

### The parasite SC is a new type of stoichiometric respirasome

To study mitochondrial SCs in parasite mitochondria, we used an intracellular pathogen model *P. marinus* from the Phylum Perkinsozoa representing an early branch of the myzozoan lineage and related to Apicomplexa, sharing morphological characteristics ^15^. To keep the complexes in a close-to-native environment and elucidate any intact SC assembly, we minimised the purification procedure. Upon mitochondria solubilisation with the mild detergent digitonin, we employed a single step of sucrose density cushion to separate the heavy components of the solubilized mitochondrial membranes. This material was subjected to native polyacrylamide gel electrophoresis (PAGE) which revealed several bands at apparent molecular weights above 720 kDa. In-gel activity assay for CIV highlighted the presence of respiratory CIV in only one assembly, the one of highest molecular weight (migrating close to 1 MDa) (Fig. S1a). The redox absorption spectrum of this sample recorded in the visible range showed presence of A-, B- and C-type hemes, characteristic of presence of respiratory complexes III and IV (Fig. S1b). The functional integrity of the CIV-containing SC was assessed polarographically using an oxygen electrode (Fig. 1a). The SC sample is active and reduces molecular oxygen at a rate of 22 ± 3 e.s^-1^ using succinate as a substrate (Fig. S1c). Addition of exogenous cytochrome *c* (Cyt *c*), but not ubiquinone, was required for activity which supports co-purification of the latter, but not of the former (Fig. 1a). The addition of the CII inhibitor malonate stops oxygen reduction and the reaction can be resumed by addition of reduced quinone to feed electrons directly to CIII (Fig. 1a). The initial III-IV rate (42 ± 13 e.s^-1^) is faster than the II-IV rate and is inhibited by the CIII inhibitor antimycin A. Subsequent addition of ascorbate and TMPD allowed recording of the CIV rate at 81 ± 31 e.s^-1^, which is sensible to potassium cyanide (KCN) (Fig. 1a, Fig. S1c). Thus, in the parasite SC electrons are transferred from CII to CIII to CIV.

**Fig. 1.**
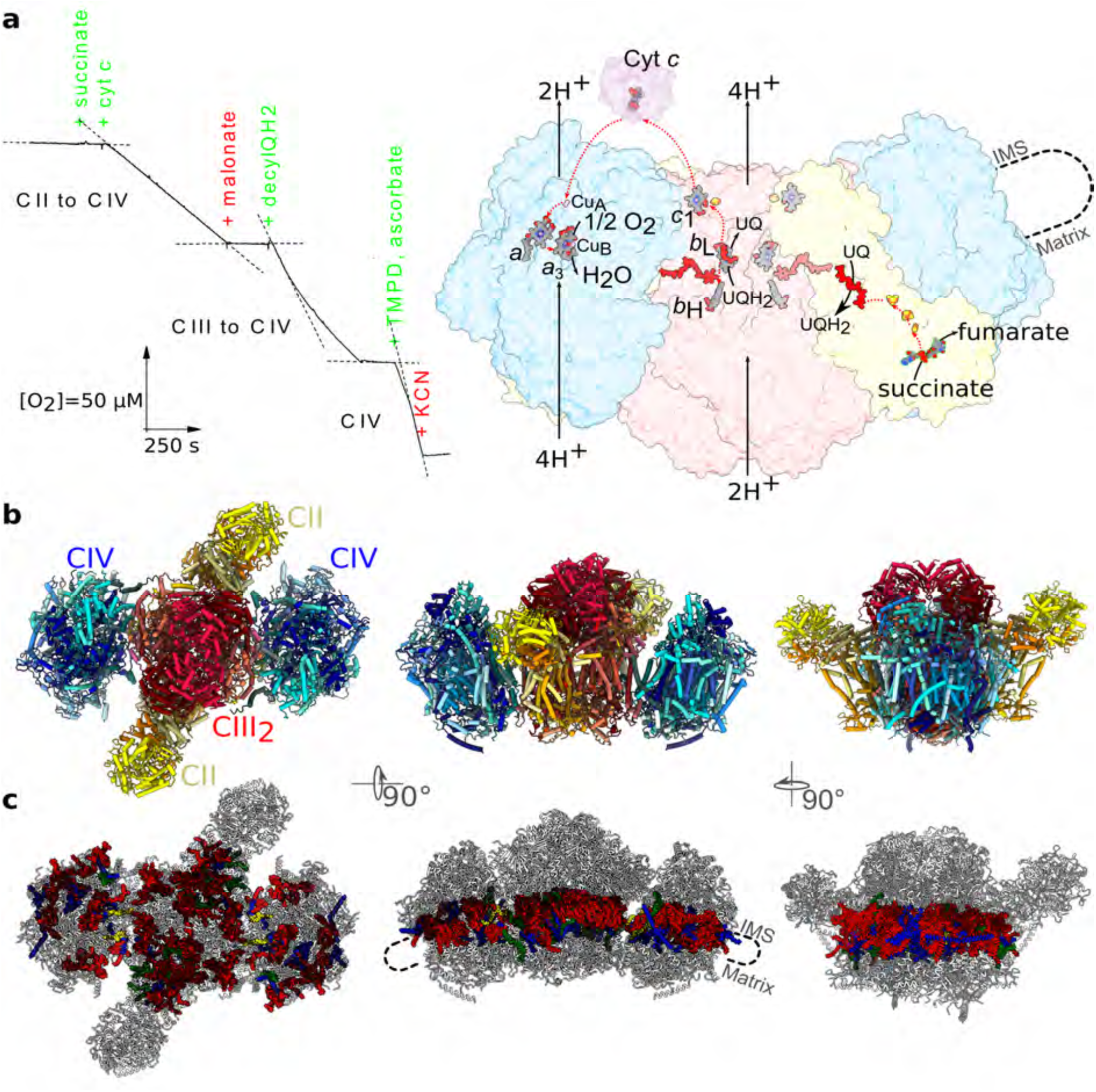
Structure of the II2-III_2_-IV2 supercomplex. **a,** Left, oxygen-consumption assay showing electron transport from succinate (via CII) and decylquinol (via CIII) to molecular oxygen (CIV). Substrates and inhibitors for the reactions are displayed in green and red, respectively. Right, Redox cofactors and electron transfer routes (red lines) underpinning the SC activity. **b,** Matrix (left) and membrane (middle and right) views of the SC model showing how CII, CIII_2_ and CIV are arranged with respect to each other. **c,** Overview of the 114 lipids modelled within the SC membrane domain, including 60 cardiolipins (red); 4 phosphatidic acids (yellow); 32 phosphoethanolamines (blue) and 18 phosphatidylcholines (green).

Since this preparation contained a mixture of high molecular weight assemblies, for structural studies, we separated them on a gel filtration column (Fig. S2a). This allowed us to reconstruct the distinctive ATP synthase hexamers (Fig. S2b) and confirm this fundamental feature of apicomplexan parasites ^12^. We then focused on the SC particles, which led to the reconstruction of the II2-III_2_-IV2 SC at an overall resolution of 2.1 Å, and the local resolution was further improved by performing masked refinements around CII2CIII_2_, CIV, and CII (Fig. S2c,d; Fig. S3; Tables S1, S3). The resulting model for the entire SC consists of 104 protein subunits, 114 lipids, and 42 co-factors for a total mass of 1.8 MDa (Fig. 1b,c; Fig. S4; Video S1, Table S3).

CII consisting of 4 conserved and 7 specific subunits (Fig. S5) is found in two copies stably bound in the SC. The presence of a single copy of CII was previously reported in ciliates ^8^, but the current arrangement in the structure of the parasite is markedly different (see below). The CIII homodimer is the most canonical of all the complexes consisting of 10 conserved and 2 specific subunits for each monomer (Fig. S6). During catalysis, the head domain of RIP1 adopts conformational changes to allow electron transfer to and from the iron-sulfur (FeS) cluster ^16^. We detected three distinct states for RIP1 in the SC and refined them to the resolution of 2.2 Å (state *b*), 2.5 Å (state *c*_1_) and 2.4 Å (intermediate) (Fig. S2d; Fig. S3b; Tables S1,S2). In the *c*_1_*-*state, the electron acceptor, Cyt *c*, docking site is occupied by a protein heterodimer that has not been previously reported and assigned directly from our cryo-EM density map to ISPR1 and ISPR2 (Fig S7). Two copies of CIV, each consisting of 12 conserved and 15 specific subunits (Fig. S8), flank the SC and appear to be more flexible. We masked and refined CIV to the local resolution of 2.2 Å (Fig. S2a, Table S1). Finally, we further studied potential conformational changes by applying a multi-body refinement analysis, which indicated continuous motion in the two eigenvectors defined by combinations of CIV rotation with respect to CIII (Fig. S9).

### Structural characterisation of the II2-III_2_-IV2 mitochondrial supercomplex

Our structure reveals an architecture of the III:IV interface that is mediated by parasite-specific subunits and influenced by the presence of CII. CIV is oriented almost perpendicular to the CIII dimer axis (Fig. 1b). Examination of the III:IV interface (Fig. 2a) shows that it is formed by six apicomplexan subunits of CIV, COX31, 33, 34, 36, 39 and COX40, and two canonical subunits, COX2B and COX6B, making the interface substantially larger than in previously reported III-IV SC structures ^17–21^. Most of the interactions formed by the apicomplexan subunits are in the intermembrane space (IMS) between a single CIV copy and one CIII monomer (Fig. 2a,b), resulting in a V-shape arrangement of the complexes that contributes to the local curvature of the membrane (Fig. S10a). Within the IMM, two single-transmembrane apicomplexan subunits, COX39 and COX40, form a helical bundle with QCR8 which is further supported by protein-lipid-protein interactions (Fig. S10b). A striking feature is that in the matrix, COX36 extends to the second CIII monomer via its subunit QCR7 (Fig. 2a). COX36 is a peripheral protein consisting of three domains: a C-terminal tail (residues 36-93) that anchors it to the CIV core in the IMS, a transmembrane α-helix (residues 10-35) that is tilted by 13° towards the second CIII monomer, and the N-terminus that forms five inter-complex interactions with QCR7 (Fig. 2d). This highlights a key role of the Apicomplexa specific COX36 subunits in tying the SC at the altered interacting surface (Fig. S11). Consequently, the overall III:IV interface includes both CIII monomers facing CIV, and a different type of SC is formed via intricate interactions of parasite-specific subunits. This is illustrated by the laterally offset CIV compared to known fungi ^17,18^, plant ^19^ and mammalian ^22,23^ SCs (Fig. S12).

**Fig. 2.**
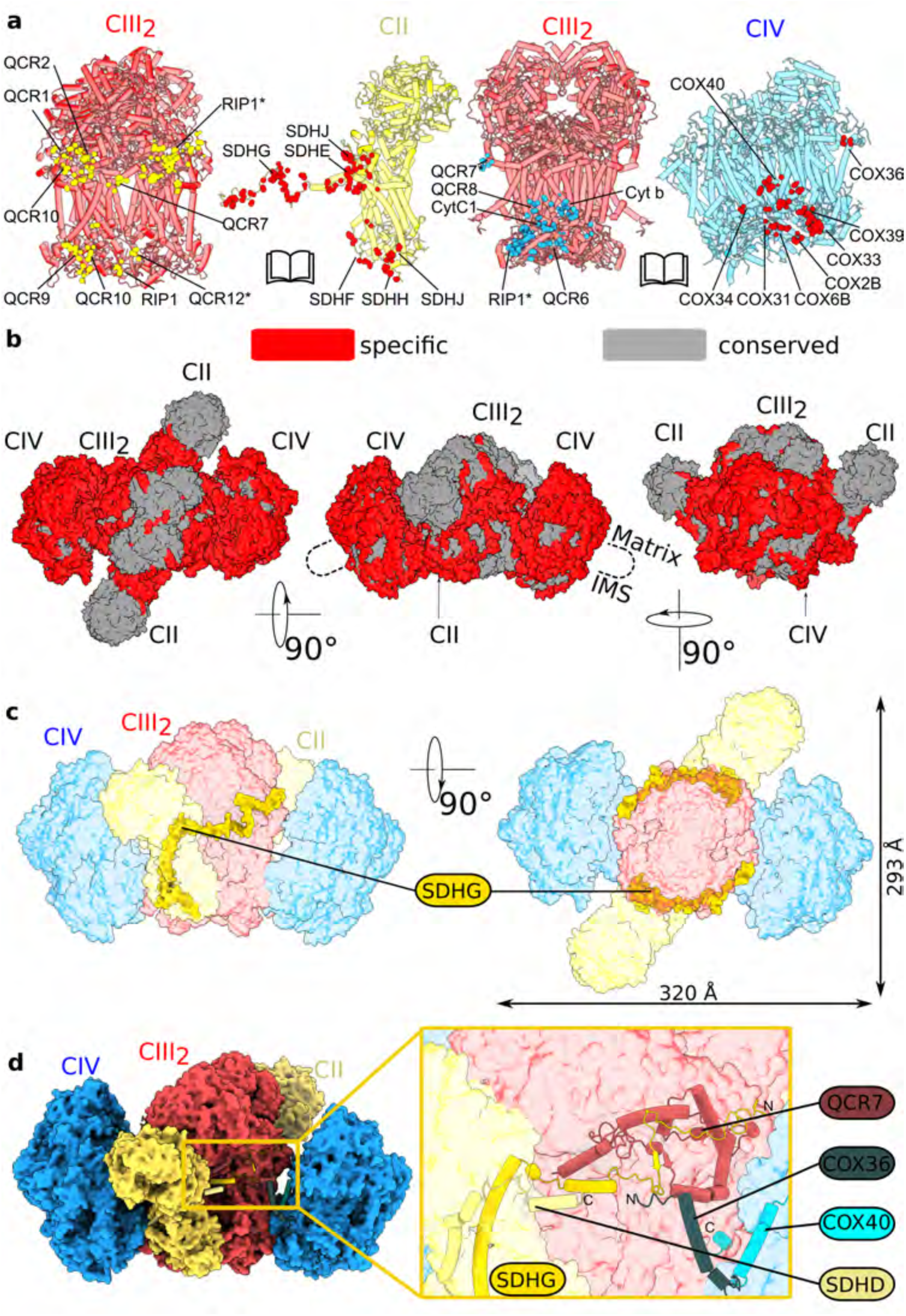
The association of CII with CIII and CIV. **a,** Open book views highlighting the contact sites of CIII_2_ with CII (left), and CIII_2_ with CIV (right). Interactions are shown as spheres (CII in yellow, CIII_2_ in red, CIV in blue). CII interacts with one CIII monomer, whereas each CIV copy interacts with both CIII monomers. **b,** The specific and conserved subunits are shown in red and grey, respectively, illustrating that the extensive contacts between complexes occur through their specific subunits, providing large interfaces in the SC. **c**, Membrane and matrix views highlighting how the N-termini of CII subunit SDHG (dark yellow) wrap around CIII_2_. **d,** SDHD and SDHG (light and dark yellow cartoon) helix bundle docks CII to CIII and CIV. Close-up view illustrating protein subunits and contact sites between the three complexes, where SDHG forms a parallel beta-sheet with QCR7 from CIII and interacts with COX36 from CIV.

The association of CII in the SC represents another feature of the structure. *P. marinus* CII has ten subunits, four conserved and six apicomplexan (Fig. S6). On the matrix side of the lipid bilayer, CII contains an amphipathic arm that protrudes towards CIII (Fig. 2c,d). The arm consists of a 13-amino acid-long helix of the apicomplexan SDHG (residues 68-80) that forms a bundle with the N-terminal extension of QCR7 (residues 4-17). Both interact with cardiolipin CDL26, highlighting the importance of co-purification of native lipids (Fig. S13). On the opposite side, it is stabilized by the C-terminal helix of SDHD, another parasite specific extension, via cardiolipin CDL19 (Fig. S13). Together, this arrangement stabilises the N-terminal region of SDHG that further wraps around CIII, engaging with RIP1 N-terminal extension over the distance of ∼45 Å (Fig. 2c). Here, SDHG (residues 49-51) also forms a parallel beta sheet with CIII subunit QCR7 (residues 28-30) and interacts with CIV subunit COX36 (N-terminal residues 1-9) (Fig. 2d, Fig. S11). Thus, all three complexes are bridged in this region of the SC.

Our structure additionally reveals that the membrane domain of *P. marinus* CII is does not hold a coordination site for a heme group as seen in other mitochondrial forms of the enzyme, including in mammals, where axial ligands are provided by the canonical subunits SDHC-SDHD. SDHC-SDHD were not previously identified in Apicomplexa due to their divergent and truncated sequences. Subunit SDHC (5.5 kDa) is shortened in our structure, representing the only protein that is found in a reduced form in the entire SC (Fig. S14a). Subunit SDHD (8.0 kDa) could only be assigned by locating topologically conserved transmembrane helices in the map. Consequently, the canonical heme-binding module adopts a different conformation, and no density for the prosthetic group could be found in our structure (Fig. S14a).

### Conformational dynamics of CIII and CIV in the supercomplex

To explore potential conformational changes relating to function, we applied multi-body refinement analysis using CIII and CIV as individual bodies (Fig. S9). The analysis revealed continuous motions describing a movement of CIV in relation to CIII (Fig. S9b). The intrinsic flexibility is defined by combinations of all rotations of CIV up to 11° (Fig. S9c). The rotations affect the distance between the electron donor and acceptor sites on CIII and CIV, respectively, where the electron carrier Cyt *c* docks. In the consensus model, the distance between the Cyt *c* docking sites on CIII and CIV is ∼79 Å (Fig. S15a). The maximum distance in the extreme states of component 1 in multi-body analysis is 84 Å and is shortened to 77 Å when maximal rotation around the pivot points is achieved (Fig. S9c). This is longer than the 60 Å measured in the yeast III-IV SC ^17,18^. Moreover, unlike in yeast, no strongly negatively charged path is observed at the surface of our SC structure that would facilitate the 2D diffusion of Cyt *c* between CIII and CIV ^24^ (Fig. S15b). Finally, in addition to the increased distance and altered surface charge, Apicomplexan subunits also form a physical obstacle for 2D diffusion (Fig. S15b). The conformational dynamics seen are primarily realised in our structure due to the COX39-COX40-QCR8 bundle at the interface, where cardiolipin CDL31 and lipid phosphate phosphatase LPP504 are resolved (Fig. S10b). The COX40 transmembrane helix assists the two complexes in maintaining the flexible membrane contacts when in the extreme conformation that reduces the distance between the donor and acceptor sites for Cyt *c* given the structural constraints. Peripheral interactions of both CIII monomers with each CIV copy further help stabilising the overall structure (Fig. 2b, Fig. S11).

### Identification of ISPR1 and ISPR2 on CIII_2_ in the *c1*-state

On quinol oxidation, electron transfer within CIII is bifurcated, and it is the *c*_1_ branch that leads to reduction of Cyt *c*. This occurs through conformational changes of the RIP1 protein head in the IMS. RIP1 is a transmembrane protein incorporated into CIII_2_ as the last maturation step to provide catalytic activity to the enzyme ^25^. In mammals, RIP1 forms a complex in the matrix with a protein derived from its own mitochondrial precursor ^26^. In our structure, the mammalian mitochondrial precursor is absent, and Apicomplexa specific protein QCR11 occupies its position, interacting with RIP1 in the matrix (Fig. S16).

To elucidate the conformational changes of the RIP1 head domain, we employed masks and classified reconstructions (Fig. S2d, S3b). This revealed three conformations of RIP1 with respect to CIII with distances between the FeS cluster and heme *c*_1_ measured as 7.4 Å (*c*_1_-state), 9.6 Å (intermediate), and 23.8 Å (*b*-state), while the corresponding distances between the UQ bound at the Qo site and the FeS cluster were 23.2 Å, 24.2 Å, and 8.1 Å. In the *c*_1_-state, comprising 38% of particles, we locally refined the rotated head area to 2.9 Å resolution, revealing additional protein density on each CIII monomers surrounding the RIP1 head domain in the IMS (Fig. S3b).

We initially constructed a polyamine backbone model into this density and estimated probable residue identities based on the map using *Coot* v0.9 ^27,28^. Predicted sequences were then queried against mass spectrometry data (Supplementary Data file), with the highest-weighted spectra displaying a sequence represented by only three unique peptides covering 25% of amino acids. We surmised that the protein database might be poorly annotated, prompting us to query the predicted sequences directly against open reading frames using BLASTP ^29^. This approach identified the shotgun sequences scf_1104296932410 1 749426 1 (https://protists.ensembl.org/Perkinsus_marinus_atcc_50983_gca_000006405/Location/View?db=;r=scf_1104296932410:1-749426) and scf_1104296971434 1 346455 1 (https://protists.ensembl.org/Perkinsus_marinus_atcc_50983_gca_000006405/Location/View?db=;r=scf_1104296971434:1-346455), which we named ISPR1 and ISPR2 due to their association with the iron-sulfur protein. The major difference between ISPR1 and ISPR2 is in their termini. ISPR1 has an extra density corresponding to 27 residues at the C-terminus, and ISPR2 has an extra density corresponding to 42 residues at the N-terminus (Fig. S7). Thus, they are present as a heterodimer of a long isoform 1 (179 amino acids) and a short isoform 2 (153 residues). The sequence of *P. marinus* (ATCC 50983) contains entries Pmar_PMAR022267 and Pmar_PMAR010955 covering 58% and 41% of the ISPR1 and ISPR2 modelled sequences with >90% identity. In addition, *Perkinsus Olseni* (ATCC PRA207) contains entries FOZ63_010970 and FOZ60_003022 covering 100% and 99% of the ISPR1 and ISPR2 modelled sequences with 65% and 80% identity. Both data have been used as reference for model building. We also found homologs for ISPR1 and ISPR2 in *Symbiodinium*, *Cyclospora*, *Toxoplasma* and *Neospora* species (Fig. S17a). TGME49_224932 from *Toxoplasma browser* has three homologous regions to ISPR, as well as an extended C-terminus (Fig. S17b). It is noteworthy that homolog search is challenging in Apicomplexa due to diverged sequences and a poorly annotated database ^30^.

### ISPR heterodimer occupies the docking site of the electron carrier Cyt *c* and locks CIII_2_ in the *c*_1_-state

Endogenous ISPR1 and ISPR2, co-purified in our structure, comprise two pairs of associated helices (α1-α2, α3-α4) separated from each other by ∼35 Å (Fig. 3a). Together, they forms two pseudo symmetrical four-helix bundles, where α1-α2 of one monomer is associated with α3-α4 of the other monomer (Fig. 3b). The *AlphaFold2* ^31^ prediction of the heterodimer using protein sequences based on the density map generates a similar secondary structure model of helical bundles, however, due to the presence of loop linkers between the helices, the ternary prediction differs slightly from our experimental model.

**Fig. 3.**
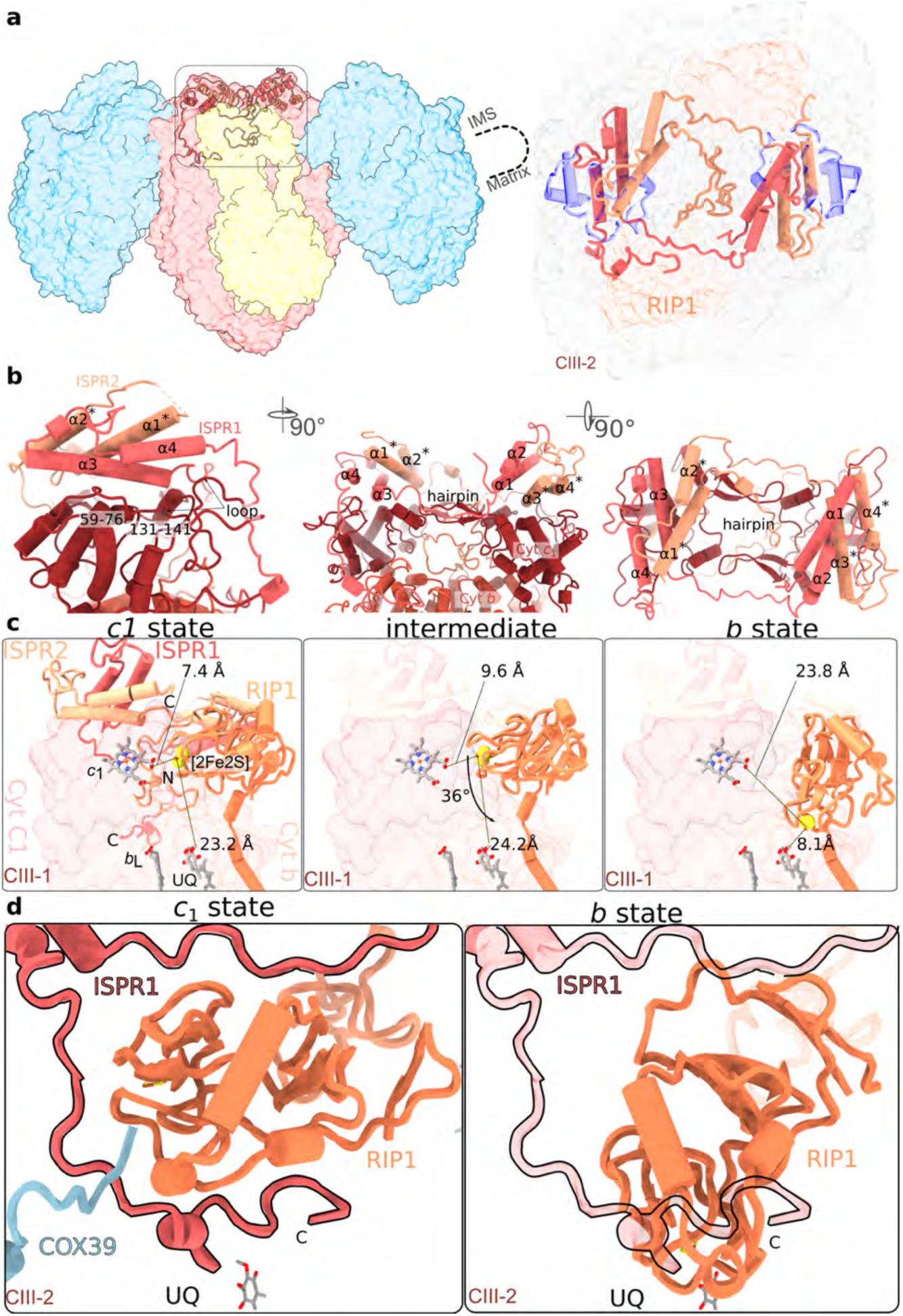
Heterodimer ISPR1, ISPR2 occupies the Cyt *c* binding site on CIII and locks it in *c*_1_ state. **a,** ISPR1 (light red) and ISPR2 (orange) in the Cyt *c* binding site. ISPR2 N-terminus fills the cavity in the IMS between the monomers of CIII, whereas ISPR1 C-terminus contacts RIP1. Superposition of the Cyt *c* (transparent purple) from the CIII_2_-bound structure (PDB ID: 1KYO) shows that the ISPR dimer compete for the same site. **b,** ISPR1 and ISPR2 contact sites on Cyt *c*_1_ (brown) and Cyt *b* (tomato). **c,** Three distinct conformations (*c*_1_ state, intermediate and *b* state) are identified by classifying the head domain of RIP1. The distances between redox centers *c*_1_, FeS, and UQ are shown. The ISPR dimer is present in the *c*_1_ state only, and its position is indicated with transparent cartoon in the intermediate and *b* state. **d**, In *c*_1_-state, ISPR1 C-terminus locks RIP head by hampering its rotation, effectively preventing electron transfer from UQ to FeS. In *b*-state, the position of ISPR1 C-terminus indicated with transparent cartoon would clash with RIP1 head domain.

ISPR1,2 binds to CIII_2_ through the solvent-exposed side of the conserved transmembrane subunits Cyt *c*_1_ and Cyt *b*. Cyt *c*_1_ provides a surface area of 1150 Å^2^ for a tight association with ISPR-α1 and α2 via four contacts (Fig. 3b): 1) loops regions 180-182 and 196-203 positioned between α1 and α2; 2) helix formed by residues 59-76 approaching the middle of α2 from the membrane plane; 3) short helix (residues 134-141) associated perpendicularly with α2 at its bending point (100°), and thus incompassed by the latter; 4) β-hairpin 105-120 that extends ∼20 Å in parallel to the α2-α3 loop (Fig. 3b). This loop leads to α3 and α4 on the opposite side that are associated on top of α1 and α2 of the other monomer, together forming the heterodimer (Fig. 3b).

In addition, ISPR1 interacts with the RIP1 head domain via residues 67-75 and the C-terminal extension (residues 138-160) running for 25 Å in the cleft between RIP1 head and Cyt *b* (Fig. 3b,c). In proximity to the ISPR1 C-terminal extension, COX39 (CIV) C-terminus engages with the head domain of RIP1. Thereby, RIP1 is locked in the *c*_1_ state (Fig. 3b,c; Video S1). ISPR2 interacts with RIP1 from the other monomer mainly via residues 105-122 and weakly via the C-terminal region, due to its shorter length compared to ISPR1. From the other side, ISPR2 N-terminus (residues 2-46) inserts into the dimer axis of Cyt *b* close to the membrane surface, aligning with the C2 symmetry of the entire SC (Fig. 3a). The N-terminal segment of ISPR2 can be further divided into two regions: the flexible linker region (residues 17-46), which traverses the central cavity of CIII_2_, and the stable axis region (residue 2-16), positioned along the C2 axis of the dimer (Fig. 3a). Thus, the ISPR1 C-terminus stabilises RIP1 head in the *c*_1_ state, and ISPR2 N-terminus further associates the heterodimer in the Cyt *c* binding site. All those interactions involve regions that are generally conserved in CIII (Fig. S6).

During the transition from state *c*_1_ to state *b*, RIP1 rotates 36^°^, so that the surface area for contact with ISPR1 moves towards UQ at the Qo site (Fig. 3c). The superposition of the two states shows that in state *b* RIP1 is positioned with its iron-sulfur head domain where ISPR1 C-terminus resides in the *c*_1_-state. Thus, the two entities would clash, and the presence of ISPR is not compatible with RIP1 in state *b*, effectively blocking the Q-cycle and preventing ubiquinol oxidation altogether. The binding of ISPR1,2 involves relatively weak electrostatic interactions compared to Cyt *c* (Fig. S15c), suggesting that the electron carrier can outcompete them for the site. The antagonistic relationship of ISPR1,2 with Cyt *c* might further reflect on regulation of electron transport, which led us to reconstitute the SC-Cyt *c* for structural characterisation, however the ISPR has not been replaced, or displaced by Cyt *c* in the study. Thus, the ISPR heterodimer is stably bound to the SC in the *c*_1_-state.

### Lipids in the supercomplex

Lipids modulate functions of membrane proteins and specifically regulate assembly and maintenance of mitochondrial SCs ^32^. Moreover, cardiolipin is involved in affecting catalytic activities of electron transfer that ultimately lead to apoptosis ^33^. A striking feature of our cryo-EM map is resolvable density for 114 lipids, and the high resolution of the reconstruction allowed identification of 60 cardiolipins, 4 phosphatidic acids, 18 phosphatidylcholines, and 32 phosphoethanolamines (Fig. 1d, Fig. 4a-c, Video S1). Overall, the lipids surrounding the II2III_2_IV2 supercomplex show curvature toward the IMS (Fig. 1d). The identification of lipids is mechanistically important because lipid molecules were shown to mediate interactions between adjacent complexes ^34,35^ and thus contribute to stabilizing the SC. Particularly, at the interaction interface of CII and CIII, in addition to previously mentioned CDL19 and CDL26, at least four other lipids are associated with subunit SDHG, mediating intercomplex interactions (Fig. 4a). In addition, at the III-IV interface, the critical interactions of QCR8 with COX39/COX40 are largely mediated by lipids (Fig. 4d). This likely provides overall flexibility to this region and explains the relative motion of CIV in our multi-body analysis (Fig. S9).

**Fig. 4.**
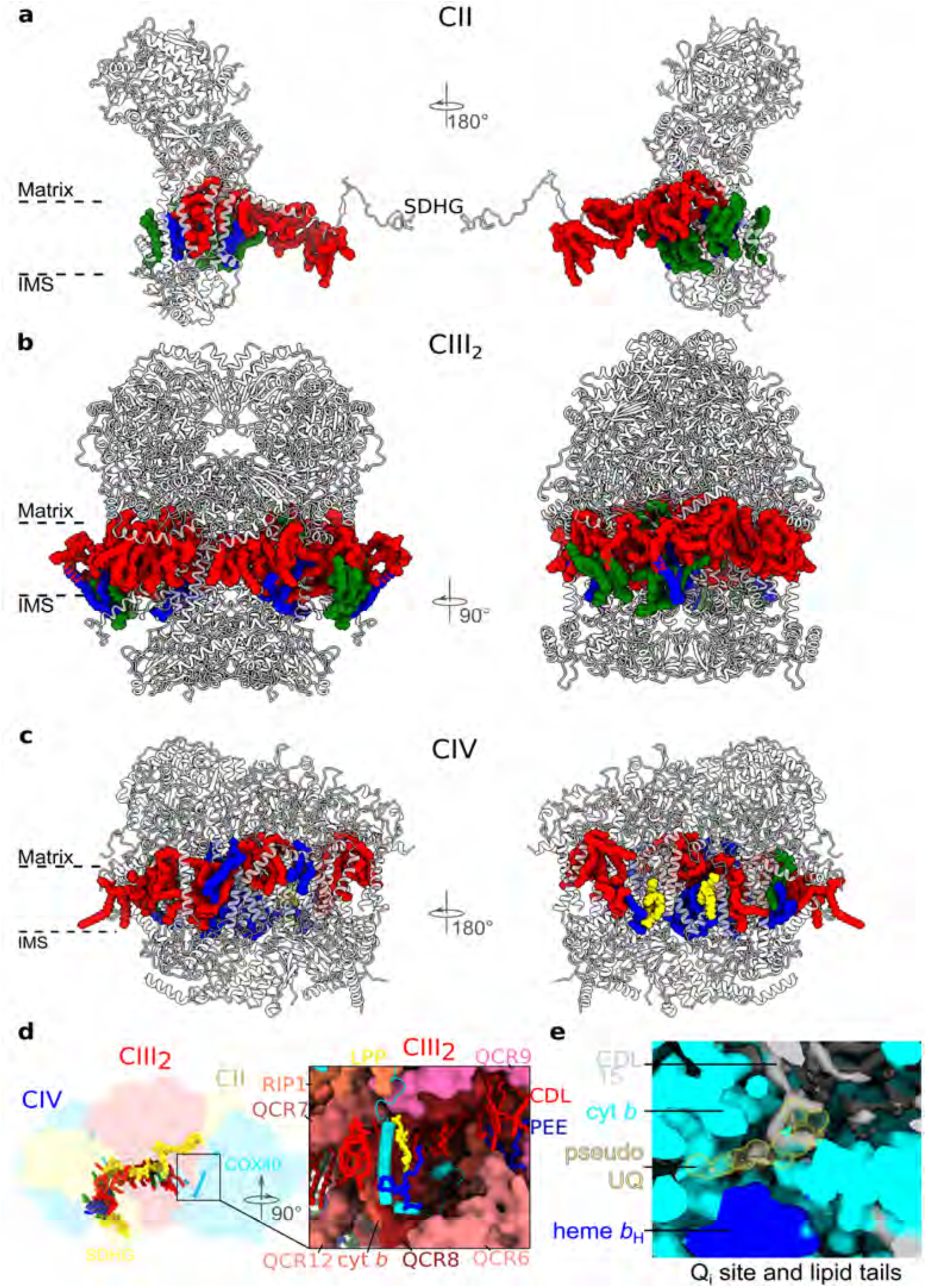
Bound resolved lipids associated with individual complexes in the supercomplex model. Two side views of CII (**a**), CIII dimer (**b**), CIV (**c**) with bound lipids. Cardiolipin (red), phosphatidylcholine (green) phosphoethanolamine (blue), phosphatidic acid (yellow). **d**, 14 specifically bound lipids to SDHG subunit of CII include 60 cardiolipins, 4 phosphatidylcholines, 32 phosphoethanolamines, 18 phosphatidic acids, which highlight their role in mediating the intercomplex interactions. Close-up view from the CIII-CIV interface further shows COX40 subunit (blue) extending into the CIII cavity (red) filled with specifically bound lipids, namely CDL8 and CDL25. During the motion of CIV related to CIII, lipids provide an environment for the COX40 extension in the cavity. **e**, In the Q_i_ site of CIII, we didn’t observe a UQ density (yellow spheres indicating potential UQ) but we observed the tail of lipid CDL15 extends here.

Lipids in our structure also refine the Q-binding pockets in CIII. Structural lipids form an architecture that facilitates Q-shuttling within CIII by two means. First, at the Q_i_ site of CIII, an elongated density is found that is not compatible with ubiquinone, suggesting that the binding pocket and the channel are preoccupied in the absence of electron carrier (Fig. 4e). Therefore, the structure suggests that the lipid would bind and unbind in coordination with the electron carrier. Further, we observed the lipid tail of CDL15 that extends to the channel of the Qi site. This provides a hydrophobic environment for ubiquinone shuttling (Fig. 4e). In addition, in CIV, we observed interactions between the farnesyl tail of heme *a* and the tail of CDL 32, supporting that the long hydrocarbon chain of heme *a* helps anchoring heme *a* into the hydrophobic environment (Fig. S18).

Finally, we show the importance of lipids in supporting the SC architecture by using *AlphaFold2* ^31^ to predict the structure of transmembrane subunits SDHG, SDHF, SDHD, SDHH, SDHE, SDHJ, SDHC, and SDHI of CII and superimposing the predicted model with our experimental structure that includes native lipids (Fig. S19). While the overall topology is generally consistent between the prediction and the experimental model, and the soluble domains fit reasonably well, most of the structural elements in the transmembrane region are shifted or rotated (Fig. S19). The differences in the local geometry are attributed to the presence of native lipids that are specifically bound and woven in between the transmembrane helices of CII (Fig. 4a). This underlies the importance of preserving the native membrane environment in structural analysis.

### *COX2* is split into three constituents with reduced hydrophobicity

The high resolution of the data allowed us to detect that the canonical mitochondria-encoded transmembrane CIV subunit 2 (COX2), which acts as the primary electron acceptor from reduced Cyt *c*, is composed of three separate polypeptides, namely COX2A, COX2B and COX2C. Remarkably, in *P. marinus* all three are encoded in the nucleus (Fig. 5a,b and Fig. S20a). Splitting of functionally important mitochondrial genes was previously reported for soluble subunits in the mitochondrial matrix ^36,37^, and for *cox2*, splitting into two genes was previously reported for dinoflagellates, alveolates, and chlorophyceae ^38^. In our structure, COX2A corresponds to the first N-terminal transmembrane helix of the mammalian counterpart; COX2B corresponds to the second transmembrane helix with 27 amino acids in the IMS that form two antiparallel beta-strands, containing the conserved residues W104 and Q194 that stabilise the histidines of the redox-active dinuclear CuA center; COX2C corresponds to the head domain that harbors the active electron acceptor site and complements COX2B with six beta-strands. Since COX2 is directly involved in electron transfer, the assembly of three proteins into the intact unit is a prerequisite for the acceptor site activation. We show that compared to the mammalian mitochondria-encoded COX2, each of the three protein segments in our structure displays a reduced hydrophobicity, resulting in an overall less hydrophobic protein (Fig. 5b). Therefore, our data supports a potential evolutionary mechanism of membrane subunit splitting end extension that would facilitate gene transfer ^39^. Together with our previous observation of a minimal nuclear-encoded subunit-a in the apicomplexan ATP synthase ^12^, the analysis illustrates how the mitochondrial genome reduction is achieved in evolution through balancing protein hydrophobicity and ensuring the correct localization in the mitochondrial membrane.

**Fig. 5.**
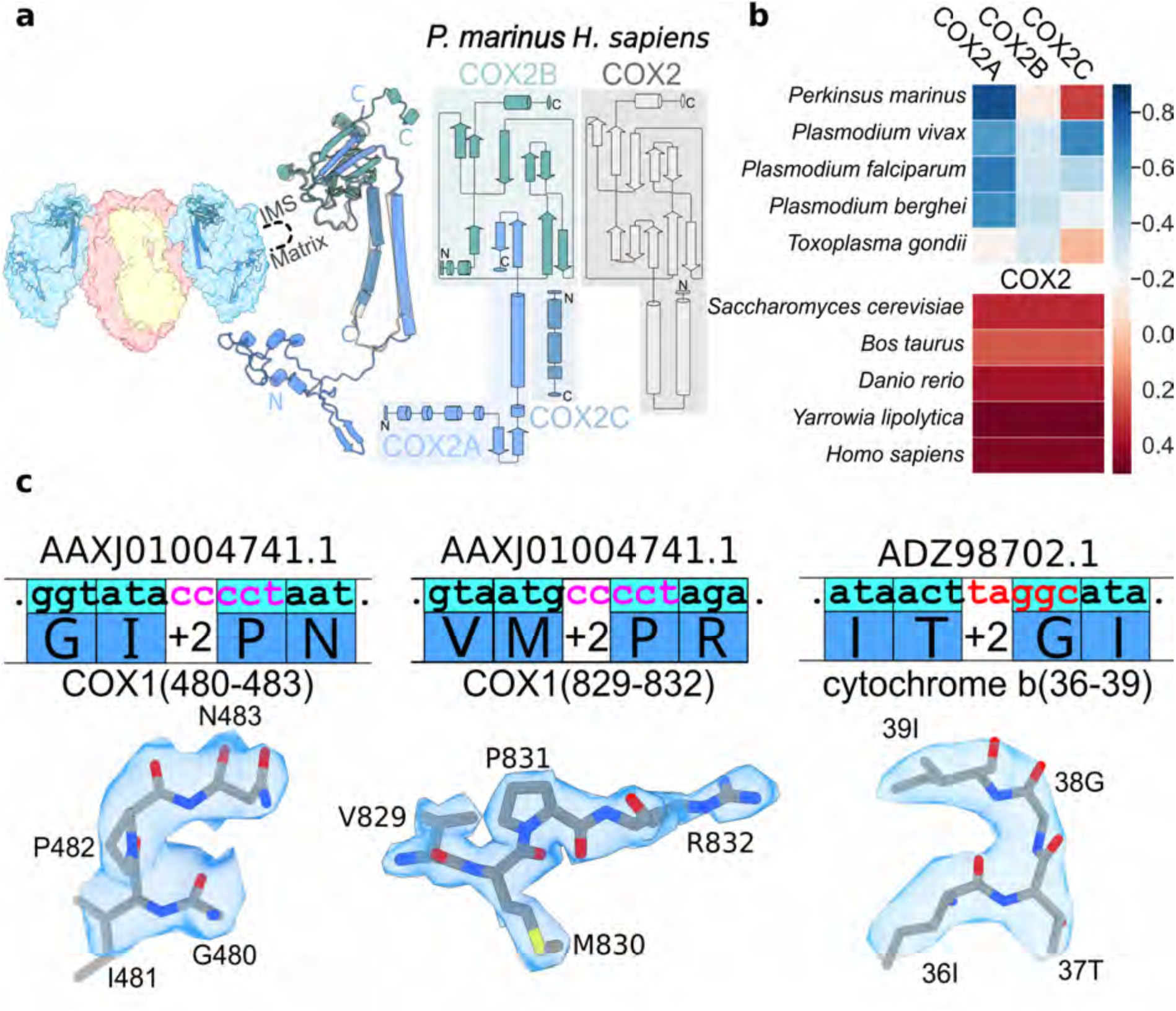
COX2 protein splitting and COX1, Cyt *b* frameshift phenomenon. **a,** Comparison of COX2 subunits of *P. marinus* and *H. sapiens* CIV. The superposition demonstrates that functional COX2 in *P. marinus* consists of three separate proteins, namely COX2A (lighter blue), COX2B (green), COX2C (darker blue) encoded in the nucleic genome. The topology diagram comparison shows the organization of the domains that are replaced by three different proteins. COX2B corresponds to the C-terminal domain, whereas a single-helix protein COX2C and one helix from COX2A collectively correspond to the N-terminal domain. **b**, Comparison of relative hydrophobicity of COX2 between apicomplexan parasites, where it is segmented into three nuclear-encoded subunits, and organisms with a single mitochondria-encoded protein. The hydrophobicity is calculated as the grand average of hydropathy according to the Kyte-J.&Doolittle(kd), Moon-Fleming (mf) or Wimley-White (ww). The nuclear-encoded COX2A, 2B and 2C of apicomplexans show a reduced hydrophobicity compared to the canonical protein. **c**, +2 frameshifts in proteins COX1 and Cyt *b* are shown, which have been identified during the model building. Original nucleotide sequence from the database is shown with the amino acids and their corresponding densities that were used to detect the frameshift.

### +2 frameshift occurs in subunits Cyt *b*, COX1 and COX3

The mitochondrial genome of the parasite is extremely reduced, coding for only three protein subunits that are Cyt *b* (CIII), COX1 (CIV) and COX3 (CIV). All three are found in our SC structure. Previous data suggested that COX1-encoding mRNA is not translated in a single reading frame with standard codon usage, but rather shifted ten times at every AGG and CCC codon by one nucleotide (+1) ^40^. This phenomenon is termed programmed frameshifting, and it enables expansion of the coding capacity of genomes and the modes through which gene expression can be regulated ^41^ A structural basis for programmed ribosomal frameshifting of one nucleotide in cytoplasmic translation has been suggested recently ^42,43^.

By comparing the modelled protein residues from our structure with the genomic sequence in the database, we detected eight frameshift events in Cyt *b*, ten in COX1, and one in COX3 (Fig. S20b). Intriguingly, three of them appear to represent switching to an alternative reading frame that is +2 nucleotides (Fig. 5c) Two are in COX1, at positions P482 and P831, and one in Cyt *b*, at position G38. To date, only an erroneous +2 frameshift has been reported in *Escherichia coli*, which occurs as a result of a translational mistake involving absence of tRNA modification ^44^ Therefore, our data provide evidence of +2 programmed translational frameshift, and likely occurs on the mitoribosome of the parasite. Our structural data are consistent with a recently updated assembly of *Perkinsus* mitochondrial genomes that identified the frameshift sites across four taxa and suggested a model where it would be driven by tRNA availability resulting in unused codons ^45^.

## Discussion

### Revisiting the ‘respirasome’ complexity: implications of CII integration

While their role remains to be fully understood, it is clear from previous studies that mitochondrial SCs exhibit compositional and structural plasticity ^1,9,46–48^. In ciliates, a stable SC containing all four respiratory chain components was shown to contribute to membrane curvature induction ^8^. Notably to date, in organisms lacking CI, only III_2_-IV2 or III_2_-IV SC structures, incapable of complete electron transport, have been reported ^17,18^. CII likely plays a substoichiometric role in optimal electron transfer, and its physical contact with the SC in CI-lacking organisms was not previously structurally implicated. However, inhibitor titration analyses suggested a CII association with CIII and CIV ^49^, leaving the possibility of direct interaction open.

To elucidate the structural and functional mechanisms of mitochondrial SCs, considering native lipids and unexplored features beyond canonical models, we determined the structure of the SC from the parasite *P. marinus* at 2.12 Å resolution. The SC is coupled to 114 native lipids and 42 co-factors (Fig. 1, Video S1). Our structure revealed two major findings: (1) CII structurally joins the SC, forming a new type of functional II2-III_2_-IV2 respirasome allowing complete electron transport from succinate to molecular oxygen; (2) a putative mitochondrial modulator, the ISPR1,2 heterodimer occupies the Cyt *c* binding site on CIII.

CII catalyses the oxidation of succinate to fumarate and increases the electrons available to CIII and CIV via coupled reduction of ubiquinone to ubiquinol in the IMM. Thus, it serves as a link between the tricarboxylic acid cycle and oxidative phosphorylation ^50^. In our structure, CII binds to CIII and CIV via specific subunits and lipids. The docking of CII occurs on the matrix side through the apicomplexan subunit SDHG, recognizing the N-terminal extension of CIII subunit QCR7 and extending to CIV subunit COX36. Structural comparisons with yeast ^17,18^ and plant ^19^ III-IV SCs shows that CII could occupy the same position as seen in the *P. marinus* SC, with no apparent predictable clashes (Fig. 6), though no proteins have thus far been identified that could mediate such interactions. The contact site between III-IV and CII may be enabled via the unoccupied membrane interface of QCR7 in those species. Thus, this new type of II2-III_2_-IV2 respirasome may not be the exclusive feature of the parasite. In a broader perspective, higher-order protein crowding in the membranes of bioenergetic organelles is an emerging pattern elucidated with recent structural studies ^1,8,12,51–53^. Our work further demonstrates that the functional arrangement of the respiratory complexes can be more intricate and diverse than previously thought.

**Fig. 6.**
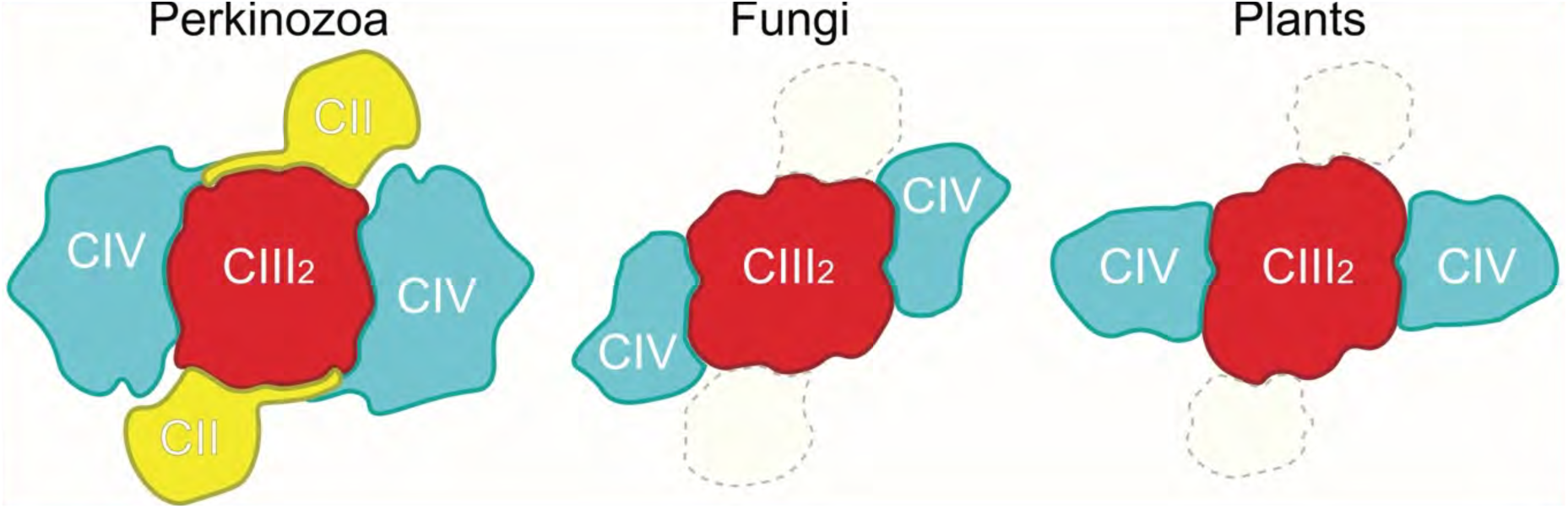
Architecture of II2-III_2_-IV2 and comparison of CII’s relative position at the fungi and plant CIII interface. Left, schematic representation of the *P. marinus*. Middle and right, fungi and plant III-IV, respectively, with their two CIV along the 2-fold axis of CIII_2_. The relative position of CII (wheat) is outlined with grey dashes, having removed apicomplexan subunits are removed from the interface.

A potential regulatory rationale of the II2-III_2_-IV2 SC association might be related to the conversion of metabolic reactive oxygen species (ROS). Approximately 90% of the cellular ROS are generated by mitochondria in the form of superoxide O2•−, which is rapidly converted to the more stable H_2_O_2_ by the enzyme superoxide dismutase (SOD) ^54^. CII and CIII are the major sites of ROS generation in mitochondria ^55^. In *Mycobacterium smegmatis*, SOD is an integral component of the SCs ^56^. In mitochondria, up to 2% of molecular oxygen is reduced to superoxide anions, and if SOD could associate similarly, the presence of CII in the SC implies proximity to SOD, facilitating a more effective defence against oxidative damage due to the abstraction of electrons directly from ROS generators during respiration. Thus, the dense organization of all the respiratory complexes would be beneficial for compartmentalization of the damaging properties.

### ISPR-mediated regulation at the CIII-Cyt *c* interface

The second major finding is that the interactions between CIII and Cyt *c* are modulated in our structure by the presence of the ISPR heterodimer. Co-purification with natural inhibitors is a common theme in the related complex ATP synthase, where its inhibitory factor 1 (IF1) enters through the open catalytic interface to mechanistically stall the central stalk rotation and ATP synthesis ^12,34,52^, which has been proposed to act as a cold-regulated switch ^57^. Our finding of the ISPR1,2 heterodimer at the highly conserved Cyt *c* binding site suggests that the activity of the SC is also regulated, and CIII is subjected to regulatory signals. Only after ISPR1,2 is removed from CIII, can electron transfer occur (Fig. 7). The deviation from symmetry between the two proteins maximizes efficiency because the ISPR1 C-terminus has evolved an extension anchoring the heterodimer to CIII, while the ISPR2 N-terminus extends to block the RIP1 head movement that is essential for enzymatic activity. Based on the weaker surface charge of ISPR1,2 compared to Cyt *c* (Fig. S15c), we tested a possible collision-type mechanism for ISPR removal by adding Cyt *c* in excess. However, ISPR1,2 was not outcompeted *in vitro*, suggesting that it is stably bound, and potentially another molecular event might trigger the removal of ISPR1,2 to facilitate enzyme activation and permit electron transfer from CIII to CIV.

**Fig. 7.**
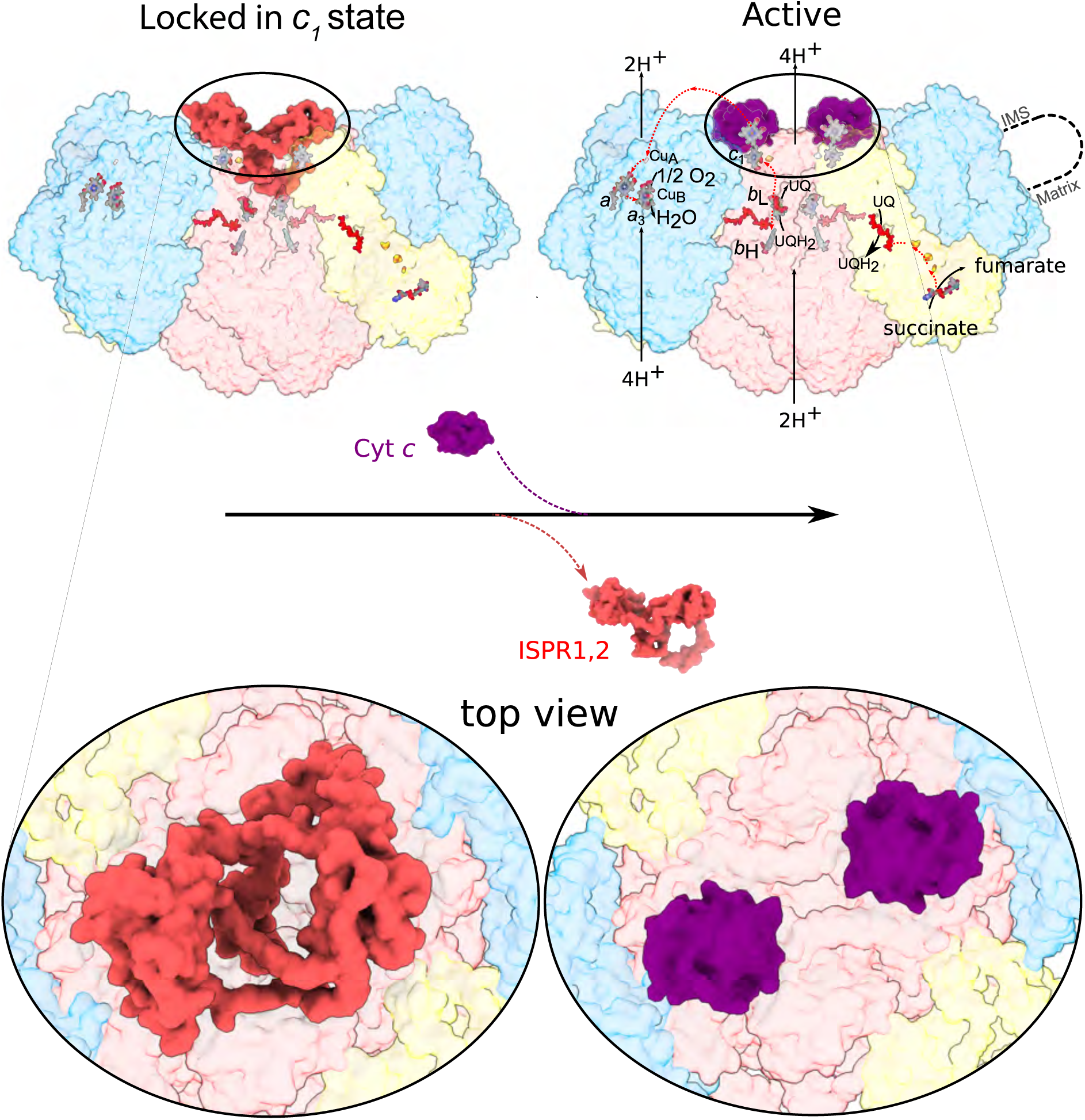
A model for SC activation. Comparison between ISPR1,2-bound (inactive) and Cyt *c*-bound structures. In the inactive state, ISPR1,2 locks CIII_2_ in *c*_1_ state. Cofactors of the electron transport chain are shown as stocks and spheres. In the active state, Cyt *c*_1_ occupies the same site (right), that permit electron transfer from CIII to CIV and enzymatic activity of the respirasome.

Biochemical and genetic studies have shown that incorporation of the iron-sulfur protein comes at a late stage of CIII biogenesis ^58^ and several proteins have been implicated in this maturation process in different species ^59–62^. However, only SCAF1, a CIV subunit isoform required for III-IV formation as well as different forms of the CI-containing respirasome in mammals ^23,45,63,64^ has been shown to remain stably bound to the mature, fully assembled CIII, on the matrix side. No chaperon or assembly factor of CIII is known in Apicomplexa. Interactions of ISPR1,2 with the RIP1 proteins clearly act as a protein inhibitor. Whether it is a regulatory factor or a facilitator of CIII or of the II2-III_2_-IV2 SC assembly, preventing premature and potentially damaging electron transfer, remains to be determined. Recently, a negative regulator of CIV has been reported in yeast, namely Mra1 that acts during the assembly of SC ^65^.

### The role of lipids in SC stability and function

In addition to the identification of new proteins in the SC, the II2-III_2_-IV2 model also contains 114 native lipids. This is important because the hierarchical organization of proteins and lipids in the bioenergetic membrane underlies the function of individual respiratory complexes and SC formation. To maintain the native lipid composition and determine key membrane components and their regulatory properties, we used a gentle solubilization and rapid purification of the material from mitochondria of parasites. The analysis of the lipids reveals their major functions: 1) dynamic interplay with electron carriers in the binding pocket; 2) lubricating role for electron carriers; 3) stabilization of intra-complex architecture; 4) contribution to inter-complex association.

With regard to the binding pocket of electron carriers, our structure contains a lipid density in the CIII active site. Therefore, the structure suggests that during the electron transfer, lipids and electron carriers continuously replace each other. This is similar to cardiolipin and cholesterol found in the c-ring of the ATP synthase from algae ^34^ and V-ATPase from synaptic vesicles ^66^, respectively, stalling its rotation, for which a dynamic mechanism has been proposed ^67^.

In addition, two structured lipids tightly bound to proteins are found around Q-binding pockets. Due to their flexible backbones, those lipids serve as solvents for Q-binding and releasing, providing a fluid film lubrication. This affects the chemical composition of the binding pocket and results in a lubricant film. The benefit of the lubrication effect is that it prevents energy losses due to decreased friction during the interfacial motion ^68^. Thus, our analysis shows that endogenous lipids have a specific shielding role in reducing activation barriers and accommodating the electron carriers. Active site lipid stabilization was also reported by high-resolution voltage-gated ion channel structures ^69^, thus it might be a more general phenomenon of membrane enzymes.

Since the resolution of the density map revealed multiple defined regular densities in the membrane space, it has allowed us to model endogenous lipids that support and stabilize the complex architecture. We illustrate this with the experimental data on CII that differs from the *AlphaFold2* ^31^ prediction in the transmembrane region, which is the critical part of the model, where some of the helices are displaced. The difference is attributed to the presence of native lipids in our structure that cannot be predicted without experimental data. This exemplifies the importance of maintaining the lipid native envelop when studying mechanisms of membrane proteins and the limitation of theoretical approaches.

Finally, the inter-complex stability in the II2-III_2_-IV2 SC is achieved due to the preparation that contains a large fraction of copurified lipids between the complexes. Out of 114 lipids, 14 and 13 are located at the II:III and III:IV interfaces, respectively, bridging the SC. This lipid composition provides a conducive environment for the assembly that can steer the formation of the SC, as previously suggested for other SCs ^22,70,71^. When the adhesive effect of lipids is maintained, it allows isolation of the higher-order SC, which is consistent with the findings from gram-negative bacteria ^72^, and bioenergetic complexes from photosynthetic systems ^35,73,74^. Thus, our structure shows that the organization of the complete respiratory chain does not only depend on the protein structure but also on the surrounding lipid environment and lipid-protein interactions. This aspect signifies the importance of the native context that must be taken into account when bioenergetic complexes are studied.

### Flexibility and evolutionary trends in SC architecture

The lipids surrounding individual complexes at the interfaces likely form a more fluid environment, fostering relative flexibility between the complexes. Comparative analysis of III-IV interface across fungi, plants, mammals, and ciliates shows species-specific protein composition with no apparent conservation (Fig. S12). This available sampling of structural studies implies a neutral evolution ^75^ potentially via mechanisms such as growing complexity with protein ratcheting ^76^, rather than a dispersal evolutionary strategy. Our structural findings suggest that in *Perkinsus* due to the additional protein acquisition: 1) no functionally important electrostatic interactions between the positively charged Cyt *c* and the SC are observed ^77^; 2) sliding of Cyt *c,* as suggested by NMR studies ^78^ and kinetic measurements ^24^, would be further hindered by the lack of a linear path and the extended distance between the sites (Fig. S15). Thus, a potential compensatory mechanism to address gratuitous protein accumulation is the conformational flexibility revealed by multi-body refinement analysis (Fig. S9). The analysis highlights how structural dynamics of the SC leads to a decrease in the distance between electron donor-acceptor sites, providing an adaptive mechanism to architectural constraints.

Two additional evolutionary aspects of the structure are the splitting of the nucleosomally-encoded COX2 subunit into three complementary segments and programmed +2 frameshifts in subunits Cyt *b* (CIII), COX1 and COX3 (CIV). For each COX2 segment, we observed reduced hydrophobicity, indicating adaptation for correct localization in the IMM. Expression of the original mitochondrially encoded proteins in the cytoplasm showed a high degradation rate and mistargeting to the endoplasmic reticulum due to recognition by the signal recognition particle ^39,79^. Thus, organelle genes encoding hydrophobic proteins are more likely to be retained in the mitochondrial genome. The segmentation of COX2 illustrates how it has evolved under selection for minimizing gene content, allowing its transfer to the nuclear genome without compromising correct localization. Selective constraint on mitochondrial gene expression that modulates its proteins is the coding capacity of the genome, which relies on tRNA availability ^45^. In this regard, our structure provides experimental evidence for such a mode of gene expression regulation, where unused codons result in a +2 programmed translational frameshift in the mitochondria of the parasite, consistent with recently reassembled mitochondrial genomes of Perkinszoa ^45^. Collectively, the evolutionary insights from our structure illustrate how a combination of the adaptive mitochondrial translation system, reduced hydrophobicity of individual gene products, and structural flexibility of the whole SC enable the function of this type of respirasome. Thus, our findings illuminate evolutionary adaptations at the gene expression and structural levels as regulatory mechanisms underlying mitochondrial genome reduction and the growing complexity of proteins in parasites.

## Conclusion

In conclusion, this work reveals the II2-III_2_-IV2 mitochondrial SC, defines ISPR as a regulator of respiration by inhibition of CIII, explains how lipids optimise the function by refining protein structure as well as ligand binding pockets, and proposes evolutionary insight into the modulation of the mitochondrial bioenergetic components. Together, it provides the structural basis for analysis of electron transport regulation and protein evolution in parasitic mitochondria.

## Methods

### Purification of *P. marinus* supercomplex

Protistan *P. marinus* was obtained from Kerafast Inc. Cells were grown at 28 ℃ until an OD_600_ of 0.6 was reached in a medium containing DMEM/F-12 (Sigma D8437), 1.82% sea salt, 0.05% glucose, 0.01% galactose, 0.01% trehalose, 1 mM Gibco GlutaMAX, 0.045 g/L cholesterol, 0.10 g/L cod liver oil fatty acids (methyl esters), 0.25 g/L polyoxyethylenesorbitan monooleate, 0.20 g/L D-α-tocopherol acetate, 2% fetal bovine serum ^80^. 14 L of media produced ∼42 gr of cell pellet. Cell pellets were washed in homogenization buffer containing 20 mM HEPES-KOH pH 7.5, 350 mM mannitol, 5 mM EDTA and lysed with glass beads (500 μm) using the FastPrep-24™ 5G Beadbeater at 6 meter/sec for 25 sec in a cold room. The lysis was assessed by a light microscope until ∼ 70% breakage was achieved. All the consequent steps were performed in 4 ^0^C or on ice. Cell debris and nuclei were separated through centrifugation at 1200 x *g* for 15 min. Mitochondria were isolated by centrifugation at 10,000 x *g* for 15 min, followed by a sucrose gradient with 15%, 23%, 32%, 60% w/v sucrose in buffer containing 20 mM HEPES KOH pH 7.5, 1 mM EDTA at 14,000 g for 60 min in SW40 rotor. Mitochondria were lysed in a buffer containing 25mM HEPES-KOH (pH 7.5), 50 mM KCl, 20mM Mg(CH3COO)_2_, 2% digitonin, 250 μL RNAase inhibitors, protease inhibitors and incubated for 30 min. The membranes were then separated by centrifugation at 30,000 x *g* for 20 min, and the supernatant was loaded on a 1 M sucrose cushion in a buffer containing 25 mM HEPES-KOH pH 7.5, 50 mM KCl, 20 mM Mg(CH3COO)_2_, 0.1 % digitonin and centrifuged at 164244 x *g* (40000 rpm) for 3 h in Ti70 rotor. Resuspended pellets were cleared by centrifugation at 30,000 x *g* for 20 min and loaded on a Superose 6 Increase 3.2/300 column. Eluted peak fractions were used immediately for cryo-grid preparation.

### Cryo-EM sample preparation, data collection and image processing

Quantifoil R2/2-300 grids floated with a homemade 3-nm amorphous carbon layer were glow-discharged before applying a 3 µl sample and vitrified in a Vitrobot Mark IV, with a 30-sec waiting time before blotting grids for 3 sec. Three data sets of a total of 43,464 movies were collected on a Titan Krios electron microscope (Thermo Fisher Scientific) operated at 300 kV using automated single-particle acquisition software (EPU) with a K3 detector at a pixel size of 0.51 Å, 0.86 Å, 0.51 Å. Data statistics are shown in Tables S1 and S2.

Movie stacks were motion corrected and dose weighted using MotionCor2 ^81^. Contrast transfer function (CTF) of the motion-corrected micrographs was estimated using Gctf ^82^. Micrographs were grouped based on optic difference, beam shift in X and Y axis using custom python scripts. ∼2,620,000 particles were picked using Gautomatch ^83^ and coordinates were imported into RELION 3.1 ^84^. Particles were extracted in a unified pixel size 0.86 Å during Bayesian polishing in RELION 3.1 ^84^, then all particles were combined and refined together. The particles were subjected to 3D classification using references generated by an initial model to separate supercomplex particles from copurified ATP synthase. The particles of supercomplex were subjected to auto-refinement to improve the angular assignments with a solvent mask applied. The obtained 3D reconstructions were used as a reference for CTF refinement. Bayesian polishing was performed, followed by another round of CTF refinement. The resulting 524,289 particles were used for auto-refinement. Masked refinements of the respective supercomplex subregions resulted in map resolutions of 2.12 Å for the entire supercomplex applying C2 symmetry, 2.16 Å for CII head, 2.17 Å for CIV, and 2.06 Å for CII2CIII_2_. Local resolution was estimated by RELION ^85,86^. The workflow is further illustrated in Fig. S2d.

Signal subtraction of CIV followed by CII2CIII_2_ refinement with a local angle search was perfomed. Focused classification using a mask around two Rieske protein was performed. The *c*_1_-state (111,717 particles), *b*-state (47,561 particles), and intermediate state (137,612 particles) were obtained. For *c*_1_ state, we further classified into two classes that are symmetry related to each other. Particles from class 2 were expanded with C2 symmetry and the unrotated particles in the expanded star file were removed to only keep one copy (class 1). Then particles of class 1 and rotated class 2 were combined and recovered back to the original particles with CIV signal and locally refined to 2.47 Å. For *b*-state and intermediate, their particles were recovered back to the original particles separately and locally refined to 2.42 Å and 2.18 Å with C2 symmetry. The results are shown in Fig. S3b.

### Model building and refinement

Models for specific proteins (Fig. S7) were built *de novo*, starting with polyalanine traces built manually in *Coot* ^28^. These initial traces were then used as templates to assign side chains or secondary structure. Protein were idnetified by comparing the assigned sequence against proteome databases. Chains 3G (g), 4Q (q), 40 (1), 2U (u), 3K (k), 30 (1) were identified using open reading frames. Detection of bulky residues W, Y, and C-C motifs was used to simplify the search space. The corresponding genes coding for those chains were eventually found in the whole genome shotgun sequences scf_110429693096, AAXJ01004741.1, AAXJ01002945.1, AAXJ01000691.1, AAXJ01000150.1, scf_1104296932410 1 749426 1 and scf_1104296971434 1 346455 1, respectively.

For chains 3G(g), 4Q(q), and 40(1), tracing sequences did not fit the density, and we manually applied frameshifts +1 or +2 to correctly fit amino acid identities to the map. Thus, all frameshifts were found independently and without using any additional information. The detected frameshifts of mitochondrial proteins COX1, COX3, and Cyt *b* with their corresponding nucleotide and protein sequences are summarized in Fig. S20, and densities for +2 frameshift events are shown in Fig. 5.

For protein sequences of chain 3L(l), we considered two hits: C5KXQ7 and C5LDF5 from protein database. C5LDF5, Aurora kinase, contains the reading frame of C5KXQ7. However, no unique peptides for C5LDF5 were identified by our mass-spectrometry analysis (see source file), thus the longer protein version has been excluded, and we assigned 3L(l) as C5KXQ7. For chain 2U, the last two residues A and M from the reading frame can’t fit the density, though the rest of residues from the translation of AAXJ01000691.1 fit the density well. Sequencing mistakes cannot be excluded.

The three states of Rieske protein head were built based in their related state maps. The lipids and digitonin were especially restrained by the cif files generated from Grade Web Server (http://grade.globalphasing.org/). Other ligands that are not in the default library were generated by eLbow in Phenix ^87^. Here, the ‘edit’ file that defines metal-coordination bonds was generated either by ReadySet in the Phenix suite or manually. Final models were further subjected to refinement with Phenix real_space_refinement v1.19.2-000 ^87^, wherein, five macro-cycles of global energy minimization with ‘edit’, secondary structure restraints, and rotamer restraints.

After one monomer was properly refined, the C2 symmetric monomer was generated from the model and the two were combined. Conflicting residues between two monomers were rebuilt in *Coot* ^28^ manually. The full model then was refined again in Phenix ^87^ with reference model restraints. The model of masked parts, CII head, CIV, and CII2CIII_2_ were extracted from the full model and refined again using related half-maps in Phenix ^87^.

Cyt *c*_1_-Cyt *c* complex in Fig. 3a was modelled with *AlphaFold2* ^31^. *Perkinsus* CII transmembrane region shown in Fig. S19 was predicted using the modified Python notebook AlphaFold2_advanced.ipynb (https://colab.research.google.com/github/sokrypton/Colab-Fold/blob/main/beta/AlphaFold2_advanced.ipynb) from ColabFold ^88^ on a local computer independent of Google Colab, where homooligomer, msa_method, and used_amber_relax are set as 1, jackhammer, and True, respectively. The following protein IDs have been used for *AlphaFold2* ^31^ modeling:

|  |  |  |  |
| --- | --- | --- | --- |
| 2O/2o | SDHG | XP_002780482.1 | C5KTH2 |
| 2P/2p | SDHF | XP_002767545.1 | C5LSM2 |
| 2Q/2q | SDHD | XP_002787086 | C5K9W0 |
| 2R/2r | SDHH | AAXJ01000252.1 (contig) | / |
| 2S/2s | SDHE | XP_002777439.1 | C5L1Y6 |
| 2T/2t | SDHJ | XP_002783522.1 | C5KJI4 |
| 2U/2u | SDHC | AAXJ01000691.1 (contig) | / |
| 2V/2v | SDHI | XP_002773883.1 | C5LCL6 |

The refinement and validation statistics are presented in Tables S1 and S2. Multiple rounds of validation and model building were carried out using MolProbity and *Coot* ^28^. For further validation, the PDB Validation server was used (https://validate-rcsb-2.wwpdb.org). The structure was analysed using *Coot* ^28^ and *ChimeraX* ^89^. Figures were prepared in ChimeraX ^89^ with additional graphical elements created using Inkscape.

### Mass spectrometry

The purified samples were lysed by 4 % SDS lysis buffer and prepared for mass spectrometry analysis using a modified version of the SP3 protein clean up and digestion protocol ^90^. 33 and 22 µg protein were alkylated with 4 mM Chloroacetamide. Sera-Mag SP3 bead mix (20 µl) was transferred into the protein sample together with 100% Acetonitrile to a final concentration of 70 %. The mix was incubated under rotation at room temperature for 18 min. The mix was placed on the magnetic rack and the supernatant was discarded, followed by two washes with 70 % ethanol and one with 100 % acetonitrile. The beads-protein mixture was reconstituted in 100 µl LysC buffer (0.5 M Urea, 50 mM HEPES pH: 7.6 and 1:50 enzyme (Trypsin) to protein ratio) and incubated overnight. Finally, the peptide samples were desalted by SCX clean up. The final peptide concentration was determined by the Bio-Rad DC Assay.An aliquot of approximately 10 µg was suspended in LC mobile phase A and 2 µg was injected on the LC-MS/MS system.

Online LC-MS was performed using a Dionex UltiMate™ 3000 RSLCnano System coupled to a Q-Exactive High Field mass spectrometer (Thermo Scientific). 3 uL was injected from each sample. Samples were trapped on a C_1_8 guard desalting column (Acclaim PepMap 100, 75 µm x 2 cm, nanoViper, C_1_8, 5 µm, 100 Å), and separated on a 50 cm long C_1_8 column (Easy spray PepMap RSLC, C_1_8, 2 µm, 100Å, 75 µmx50cm). The nano capillary solvent A was 95% water, 5%DMSO, 0.1% formic acid; and solvent B was 5% water, 5% DMSO, 95% acetonitrile, 0.1% formic acid. At a constant flow of 0.25 μl min^-1^, the curved gradient went from 6%B up to 43%B in 180 min, followed by a steep increase to 100%B in 5 min.

FTMS master scans with 60,000 resolution (and mass range 300-1500 m/z) were followed by data-dependent MS/MS (30 000 resolution) on the top 5 ions using higher energy collision dissociation (HCD) at 30% normalized collision energy. Precursors were isolated with a 1,2m/z window. Automatic gain control (AGC) targets were 1e6 for MS1 and 1e5 for MS2. Maximum injection times were 100 msec for MS1 and MS2. The entire duty cycle lasted ∼2.5 sec. Dynamic exclusion was used with 60 sec duration. Precursors with unassigned charge state or charge state 1 were excluded. An underfill ratio of 1% was used.

The MS raw files were searched using Sequest-Percolator or Target Decoy PSM Validator under the software platform Proteome Discoverer 1.4 (Thermo Scientific) against *P. marinus* (strain ATCC 50983) Uniprot database and filtered to a 1% FDR cut off.

We used a precursor ion mass tolerance of 10 ppm, and production mass tolerances of 0.02 Da for HCD-FTMS. The algorithm considered tryptic peptides with a maximum of 2 missed cleavage; carbamidomethylation (C) as fixed modification and oxidation (M) as variable modifications.

### Gel Electrophoresis, protein staining and in gel-activity assay

Pre-cast gels (NativePAGE™ 3 to 12%, BIS-TRIS, 1.0 mm, Mini Protein Gel, Invitrogen) were used for Clear Native (CN) PAGE with two identical sample lanes of the purified super-complex sample post sucrose cushion and a protein ladder lane (NativeMark™, Invitrogen), prepared according to the manufacturer’s instructions. The gel electrophoresis was conducted at 4°C, firstly at 150 V for 30 min with NativePage Light Blue Cathode buffer and then at 250 V for 150 min with NativePAGE Anode Buffer. The two sample lanes were then cut. One was stained with Coomassie Brilliant Blue overnight and then destained with an ethanol/acetic acid solution. The other sample lane was incubated for 45 min in a solution of 50 mM KPi pH 7.4 containing 0.5 mg/mL 3,3′-diaminobenzidine (DAB) and 1 mg/mL equine cytochrome *c* before incubation in 10 % acetic acid.

### Activity measurements

The functional integrity of the SC was assessed polarographically using an oxygen electrode (Oxygraph+ system, Hansatech Instruments). All measurements were carried out at 25°C in a solution of 10 mM KPi, 50 mM KCl at pH 6.6 supplemented with 500 units/mL superoxide dismutase, 250 units/mL catalase and 0.05% digitonin. A typical experiment started with addition of 20-30 nM SC and the rate left to stabilise before addition of 10 mM Na^+^-succinate (substrate of CII) followed by 50 μM equine cytochrome *c*. The CII-to-CIV reaction was stopped with addition of 10 mM Na^+^-malonate (inhibitor of CII). Subsequently, 40 μM decylubiquinol (substrate of CIII) was added before the CIII-to-CIV reaction was stopped with addition of 10 μM antimycin A (inhibitor of CIII). Finally, 2 mM Na^+^-ascorbate and 40 μM TMPD were added and the CIV reaction thereafter stopped with addition of 1 mM KCN. The reaction vessel was washed with a solution containing BSA in between experiments to remove any residual of inhibitors.

Reaction rates were derived from the oxygen consumption rate recorded after addition of each specific substrate using the final KCN-inhibited rate as a baseline. Conversion to turnover numbers was made using the concentration of CIV in the sample, assuming that in the purified SC sample, all the CIV is in the SC, as supported by in-gel activity performed on CNPAGE gel. CIV concentration was determined from a dithionite-reduced *minus* oxidised difference visible spectrum recorded in 25mM HEPES, 25 mM KCl, 5 mM MgCl_2_ and 0.1% digitonin at pH 7.5 using the specific difference in absorption associated to the reduction of the A-type hemes of CIV (ΔA 603-621 nm, extinction coefficient of 26,000 M^-1^cm^-1^).

### Data availability statement

The atomic coordinates were deposited in the Protein Data Bank (PDB) under accession numbers XXXX (*c*_1_ state), XXXX (*b* state), XXXX (intermediate). The cryo-EM maps have been deposited in the Electron Microscopy Data Bank (EMDB) under the respective accession numbers: EMD-1XXXX *c*_1_ state combined map), EMD-1XXXX(*b* state combined map), EMD-1XXXX (intermediate combined map), EMD-1XXXX (*c*_1_ state CII_2_CIII2), EMD-1XXXX (*b* sate CII2CIII_2_), EMD-1XXXX (intermediate CII2CIII2), EMD-50354 (concensus II2III_2_IV2), EMD-1XXXX (CII head), and EMD-1XXXX (CIV) and EMD-1XXXX (CII2CIII_2_). An Excel file containing the visible absorption spectroscopy data and their analysis has been added. All the data will be publicly available. Source data are provided with this paper.

## Acknowledgments

We thank the European Research Council (ERC-2018-StG-805230 to A.A.), the Medical Research Council UK (MR/T032154/1 to A. Maréchal). The cryo-EM facility is funded by the Knut and Alice Wallenberg, Family Erling Persson, and Kempe foundations.

## Author contributions

A. Mühleip, L.S. and A.A. designed the project. F.W. and A.A. performed cell culturing, isolation of mitochondria and prepared the sample. F.W. collected and processed cryo-EM data and built the model. T.G., A. Maréchal and A.A. performed biochemical and spectroscopic analyses. F.W., A. Maréchal and A.A. wrote the manuscript.

## SUPPLEMENTARY INFORMATION

**Supplementary Fig. 1:**
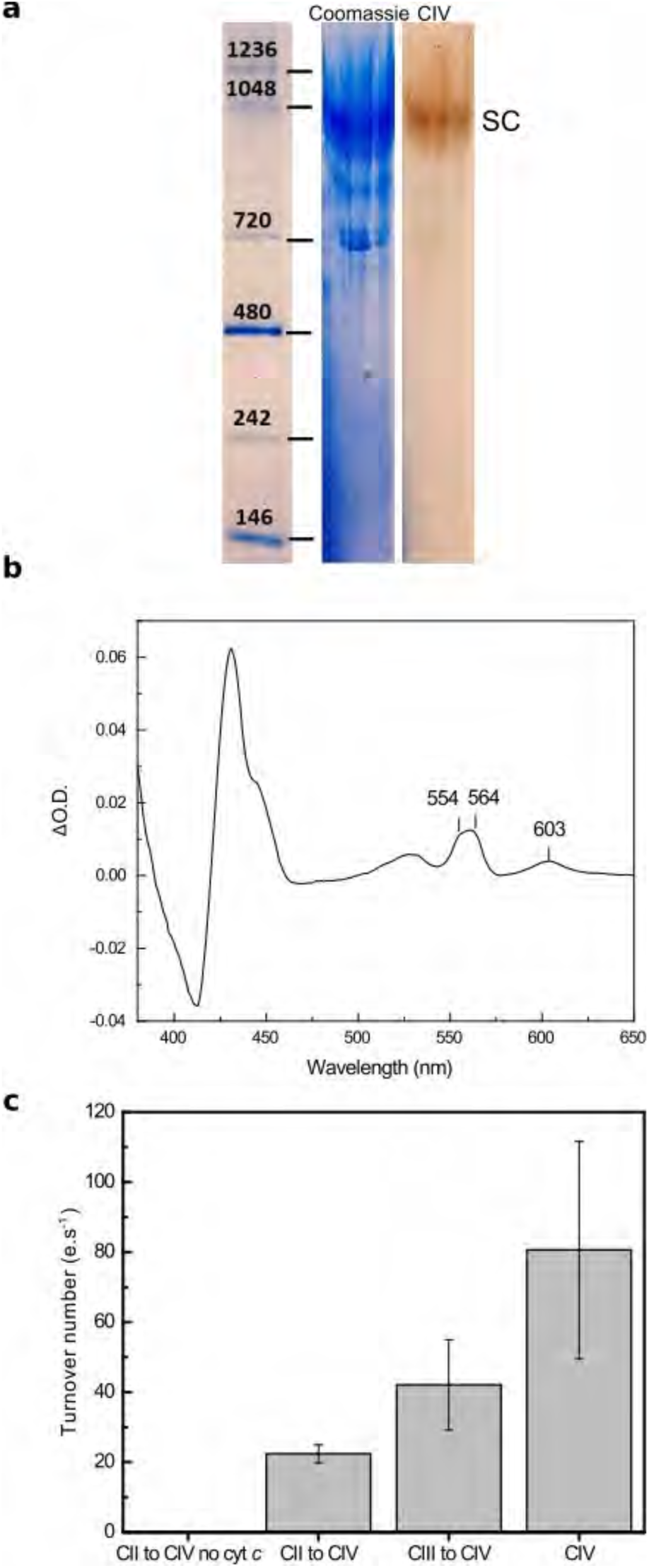
Purification and biochemical characterization of the respirasome from *P. marinus*. **a.** Redox visible absorption spectrum of the SC sample obtained after sucrose cushion revealing the presence of C- (554 nm), B- (564 nm) and A- (603 nm) type hemes, characteristic for CIII and CIV. **b.** Turnover number (*e.s^−^*^1^) measured for the SC on oxygen consumption with substrate succinate (CII to CIV no cyt *c*), with succinate and additional cyt *c* (CII to CIV), with substrate decyl QH_2_ and cyt *c* (CIII to CIV), with cyt *c* and ascorbate and TMPD (CIV). Values given are means ± standard deviations.

**Supplementary Fig. 2:**
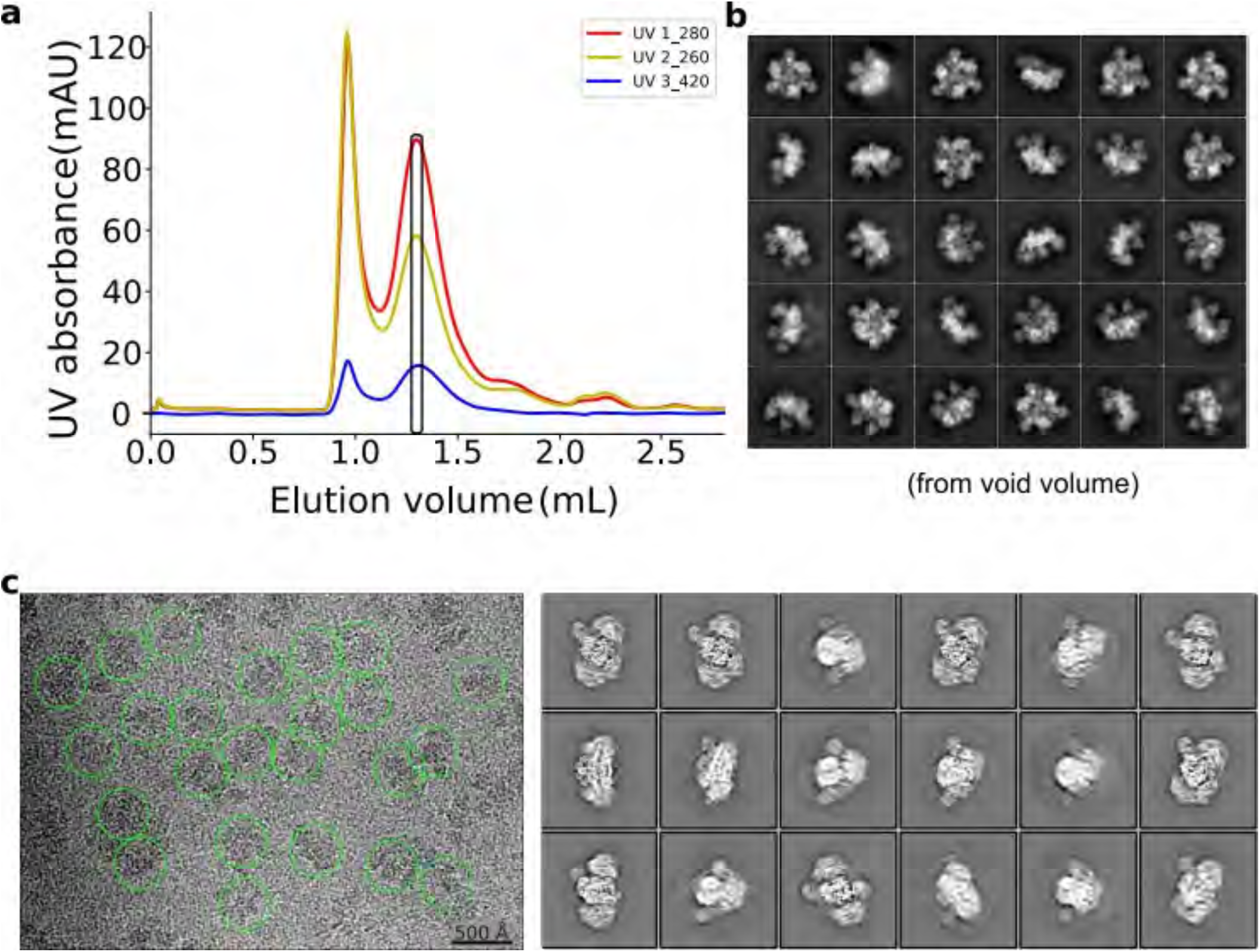
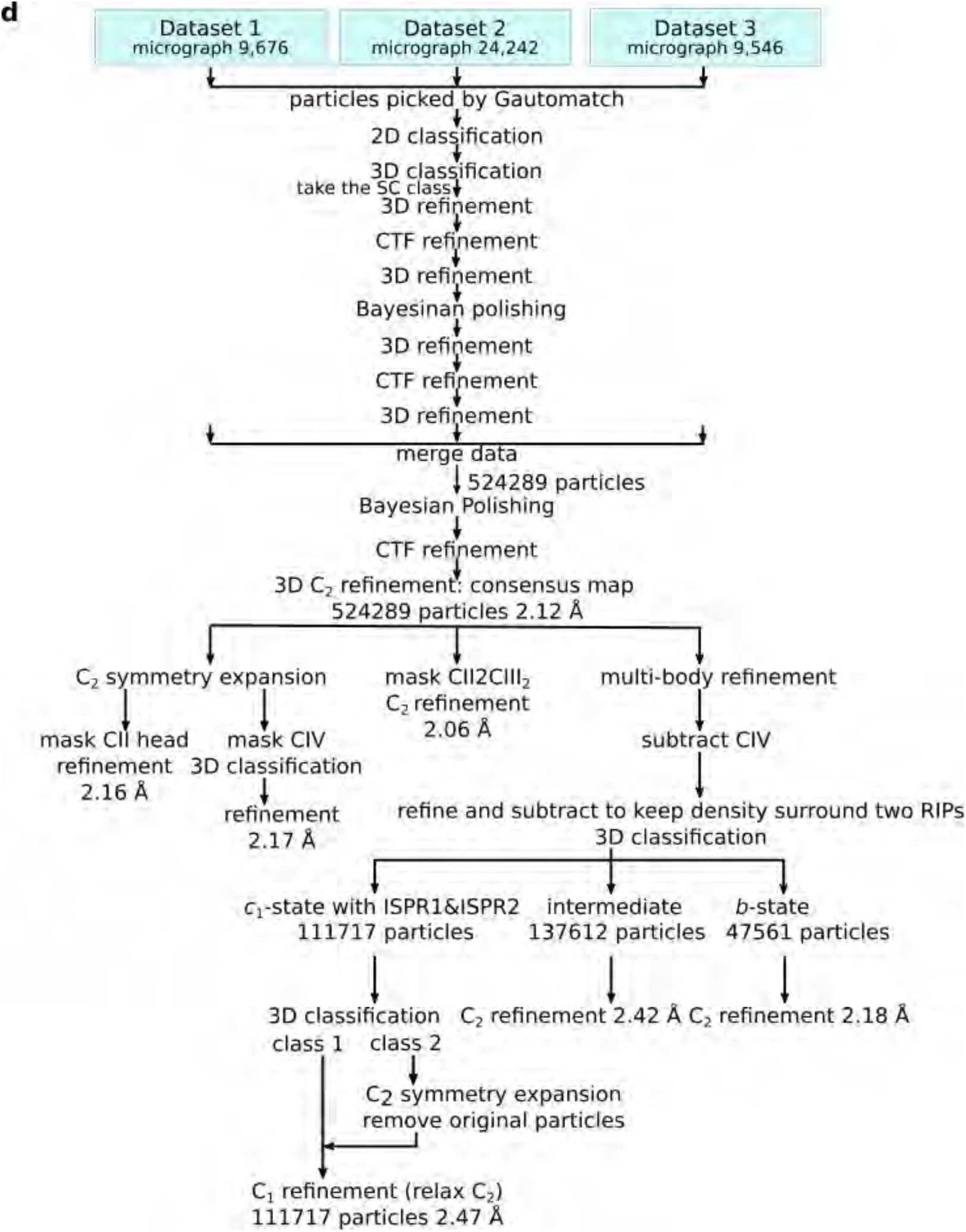
Cryo-EM image processing overview. **a.** Chromatogram of the Superose 6 Increase 3.2/300 column, the pooled fractions used for cryo-EM are indicated with a rectangle. **b.** 2D class averages of ATP synthase hexamers from void volume. **c.** A representative micrograph with picked particles (green circles). 2D class averages of SC. **d.** Data processing scheme for the complete SC II2-CIII_2_-IV2, CII head, CIV, CII2CIII_2_, and CIII states *c*_1_, *b*, and intermediate. A total of 43464 movies were recorded and analyzed, particles that were not SC were discarded by classification, since they cannot contribute to reconstruction. Consensus map was produced from 524289 particles. Cryo-EM structures were successfully obtained from three preliminary datasets.

**Supplementary Fig. 3:**
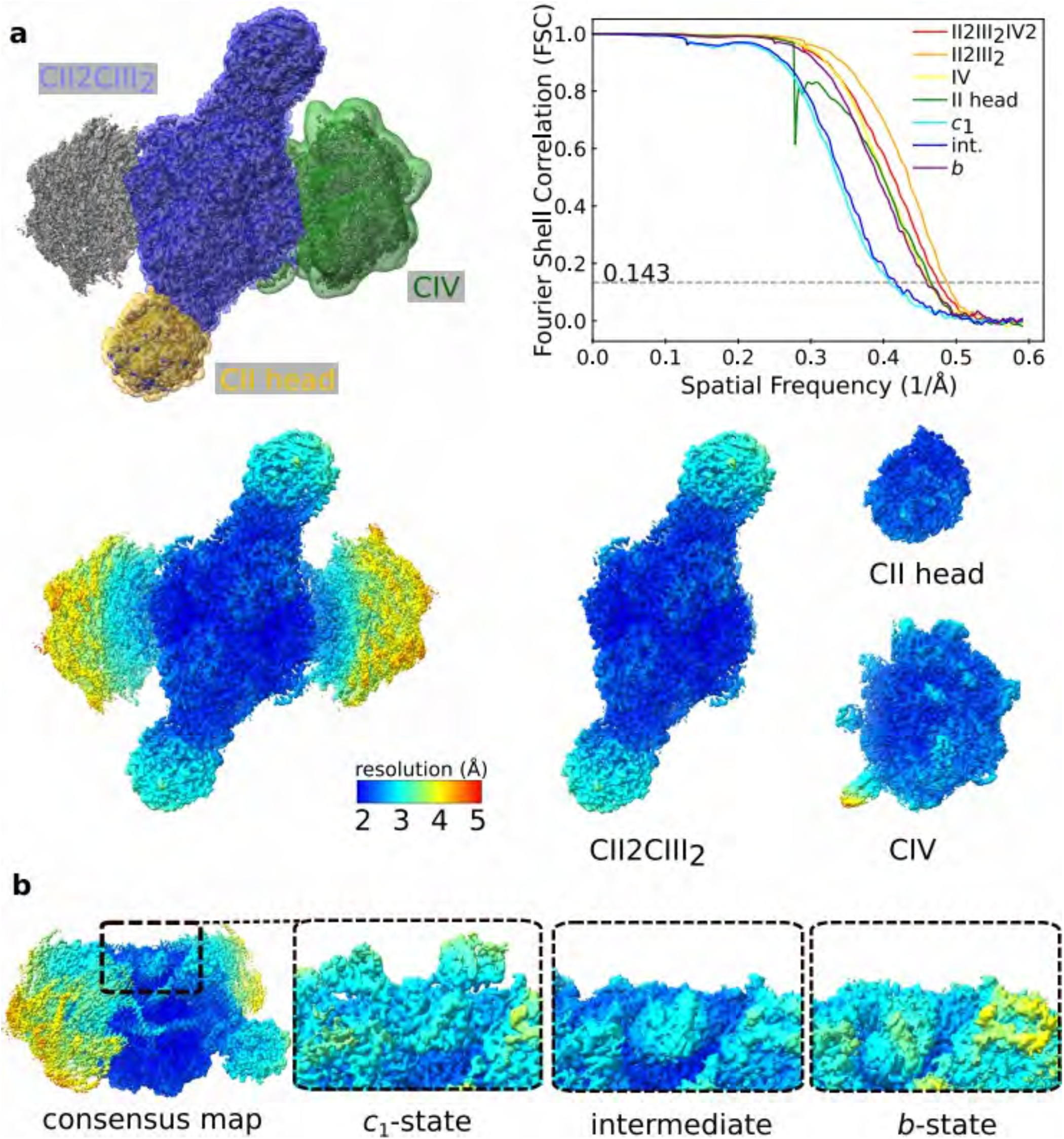
Map quality for individual complexes and states. **a.** Binary masks used for local masked refinements of CII2CIII2, CIV, and CII head. Fourier Shell Correlation curves of the half maps and local-masked refinements. Cryo-EM map resolution estimates by Fourier Shell Correlation were performed using half maps from random half-sets. Consensus map colored by local resolution. Local-masked refinements colored by local resolution. **b.** Local resolution of maps for RIP1 when in *c*_1_,*b*, and intermediate states.

**Supplementary Fig. 4:**
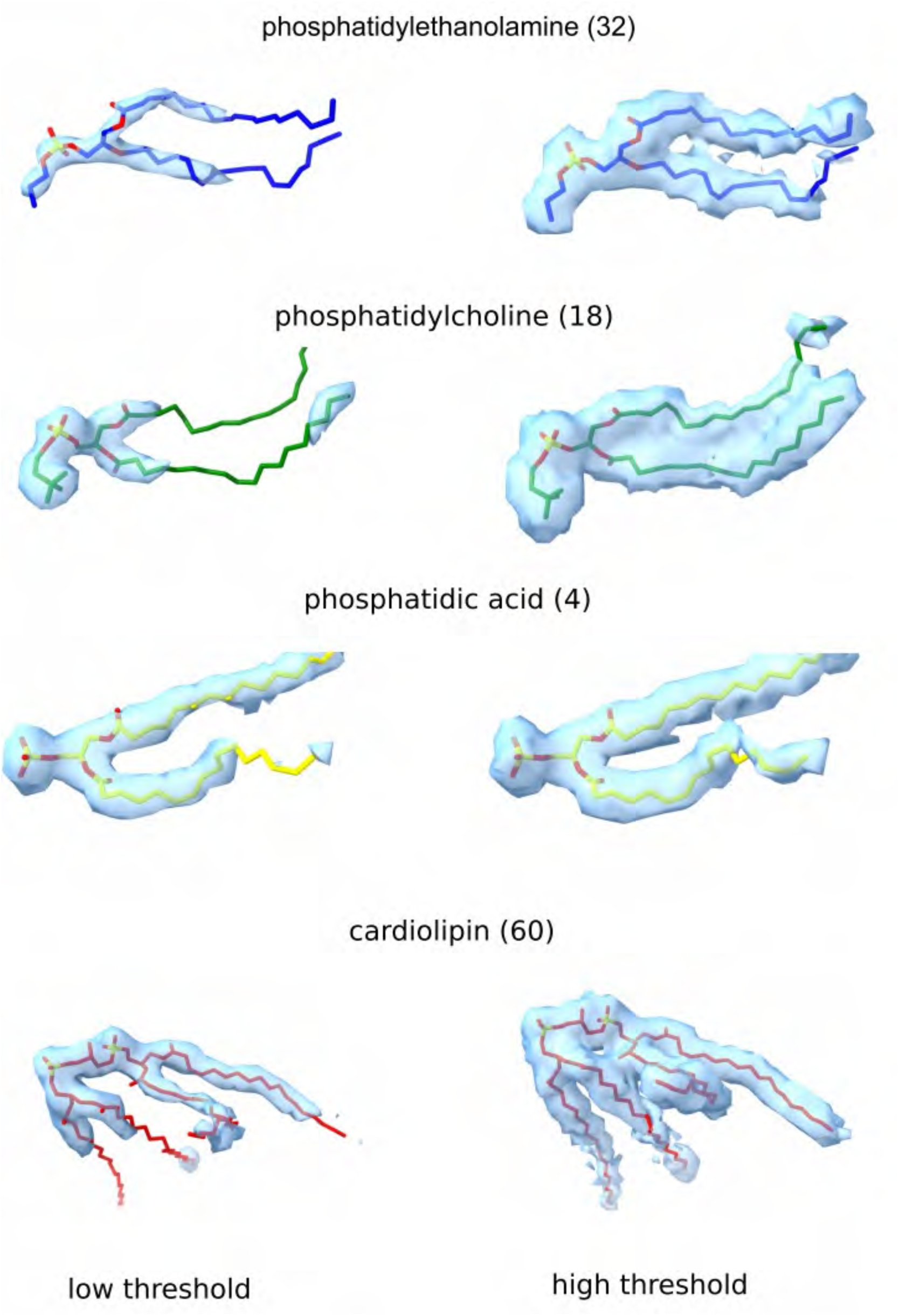
Endogenous structured lipids identified in the map. Four types of lipids are shown with their densities in two different thresholds. For each type, the number of modeled lipids is indicated in the brackets.

**Supplementary Fig. 5:**
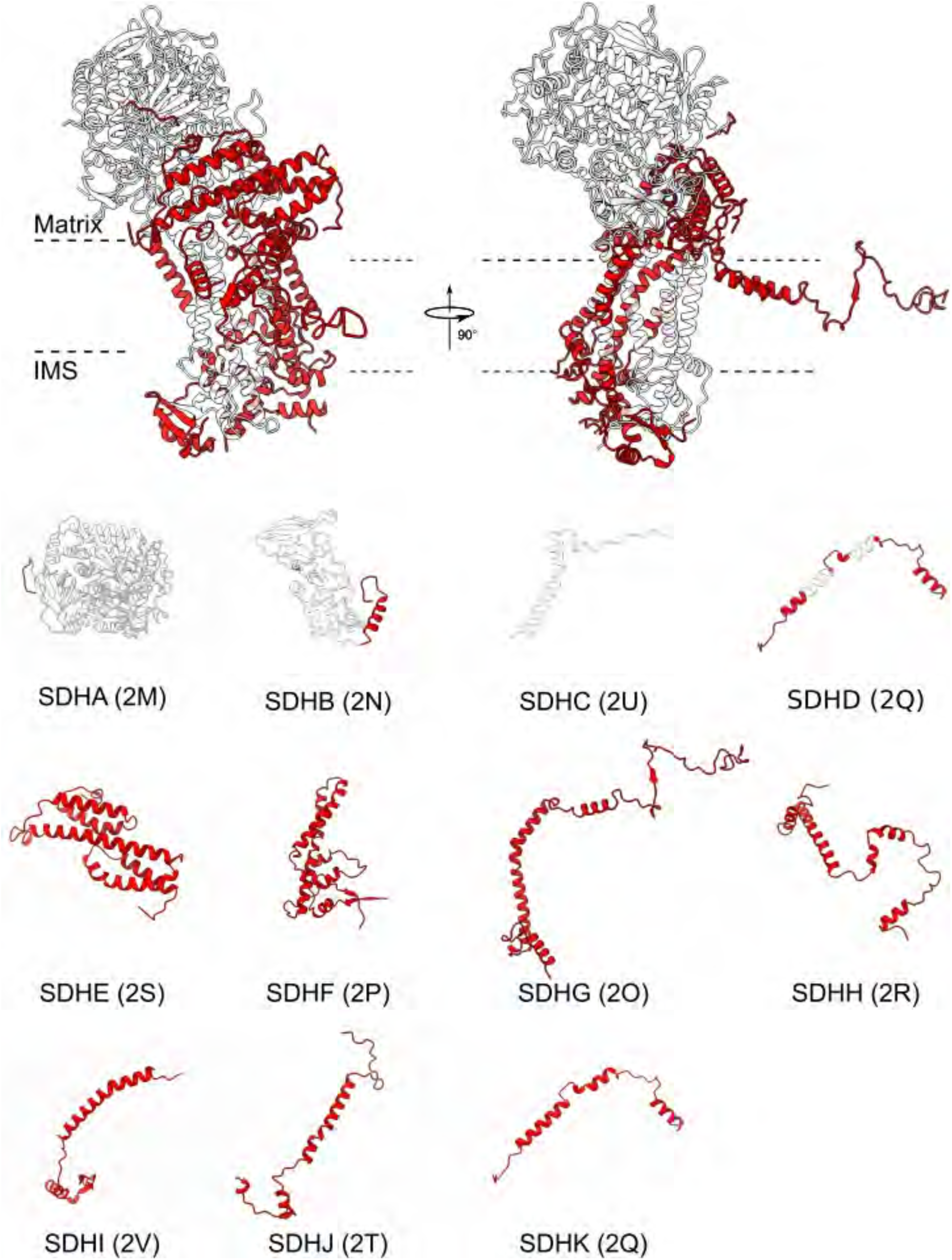
Cartoon representation of the individual subunits forming CII. Tertiary folds of all the built components are shown. Conserved structures with the canonical (mammalian) CII are in white; extensions and specific proteins are in red. Chain IDs are indicated.

**Supplementary Fig. 6:**
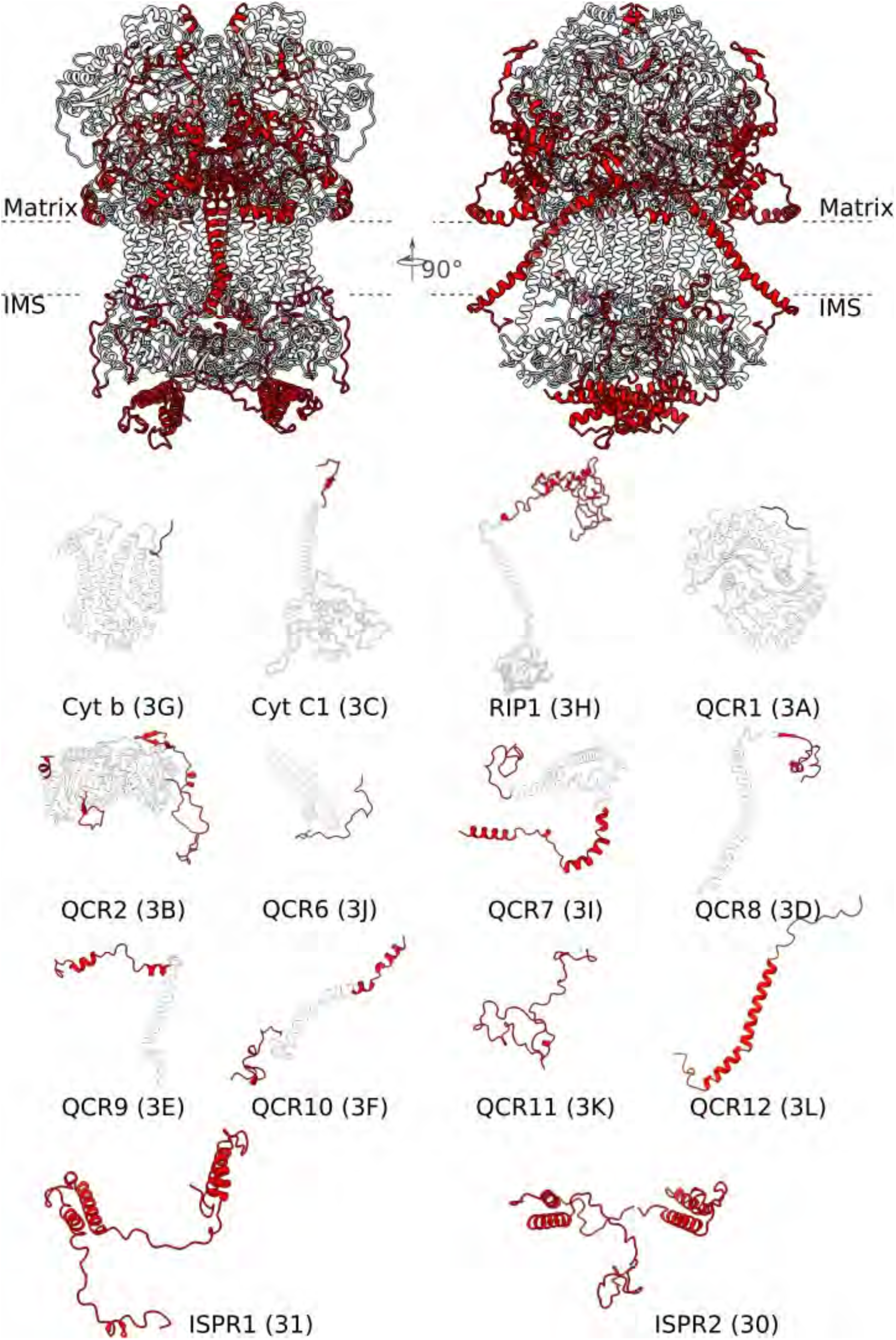
Cartoon representation of the individual subunits and associated proteins of CIII. Tertiary folds of all the built components are shown. Conserved structures with the canonical (mammalian) CIII are in white; extensions and specific proteins are in red. Chain IDs are indicated.

**Supplementary Fig. 7:**
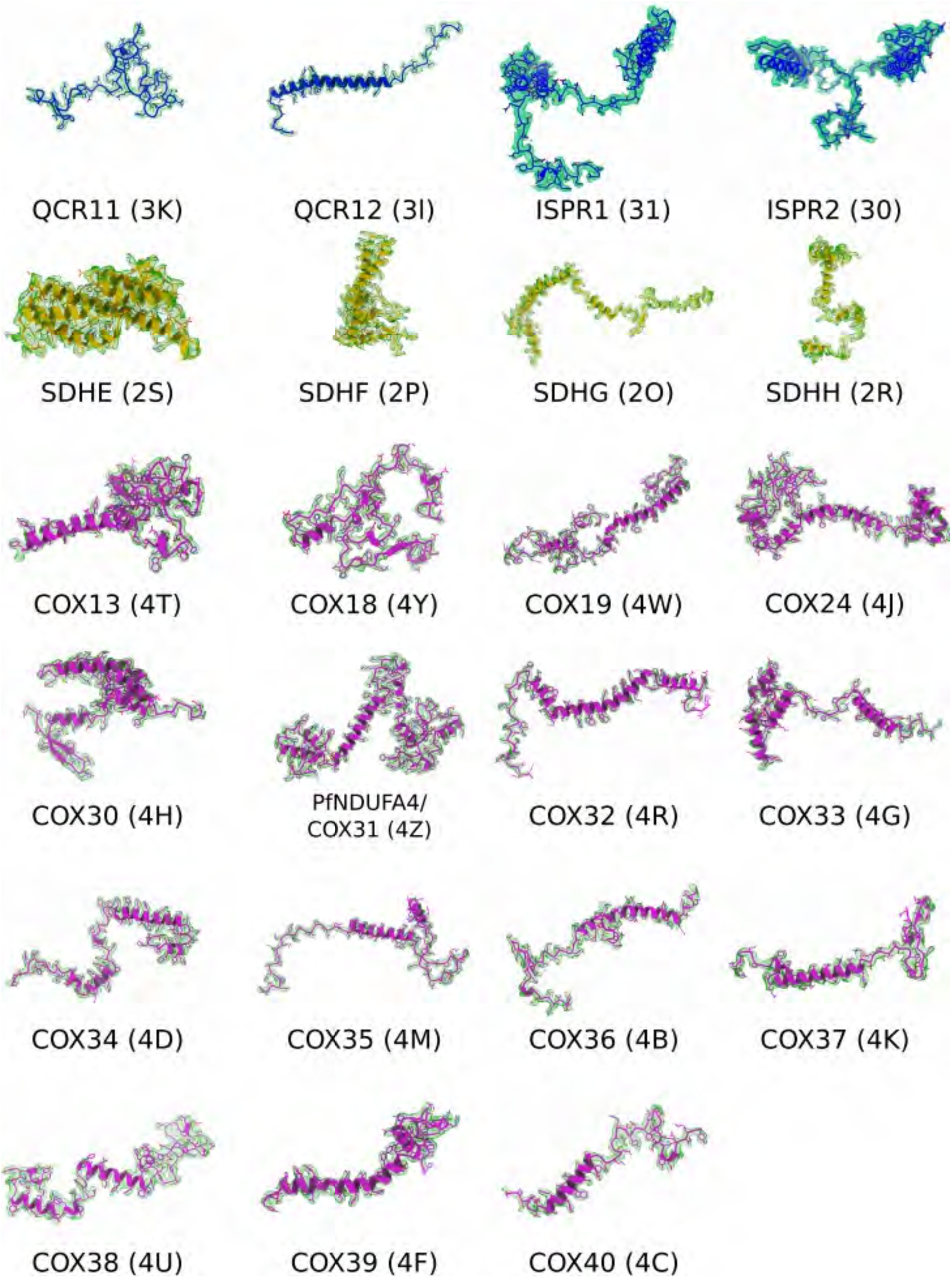
Newly modeled subunits in the SC. Specific subunits identified directly from the complex in which they associate. CII yellow, CIII blue, CIV magenta. The ISPR1 and ISPR2 heterodimer occupies the Cyt *c* electron acceptor binding site.

**Supplementary Fig. 8:**
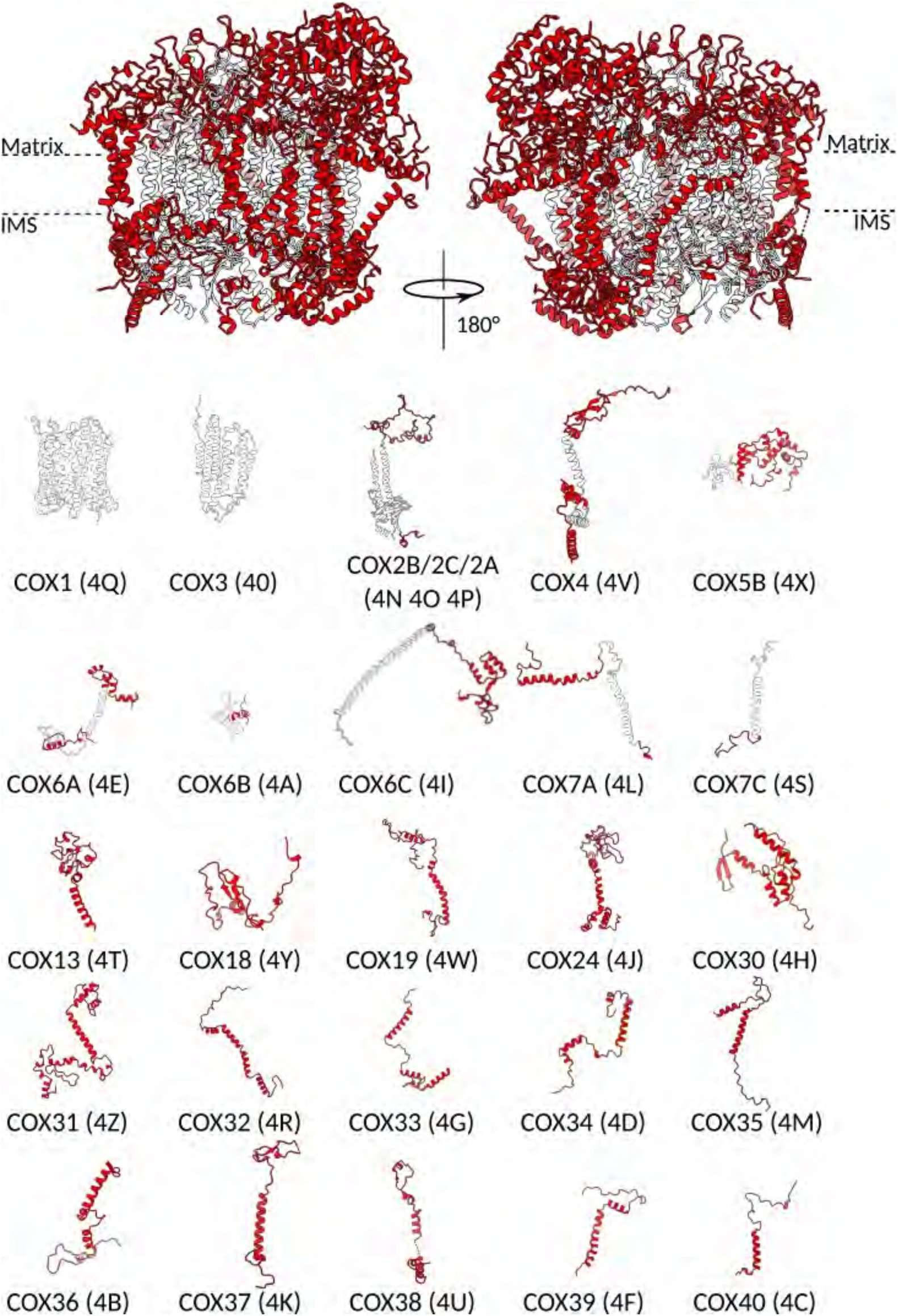
Cartoon representation of the individual subunits forming CIV. Tertiary folds of all the built components are shown. Conserved structures with the canonical (mammalian) CIV are in white; extensions and specific proteins are in red. Chain IDs are indicated.

**Supplementary Fig. 9:**
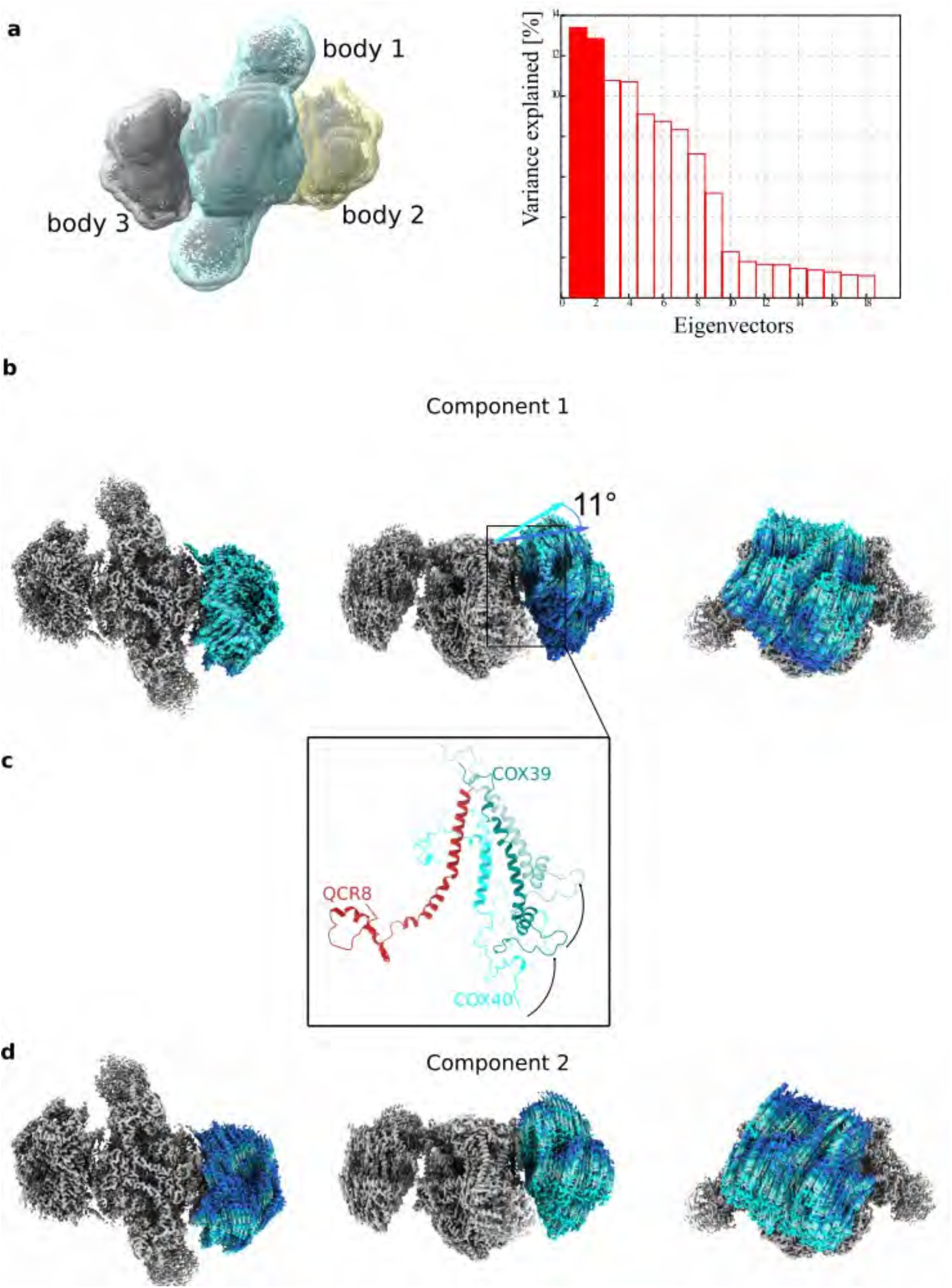
Multi-body refinement analysis. **a.** Three masks used for multi-body refinement: body 1 – CIII_2_, body 2 – CIV(1), body 3 – CIV(2). Eigenvectors that explain the variability of the data. The distribution of eigenvectors shows that none of them accounts for more than 14% of the motion. **b.** Top, front, and side views of the map, showing the motion along the component 1. **c.** The two extreme positions of COX39 and COX40 in relation to fixed QCR8 in component 1. **d.** Top, front, and side views of the map, showing the motion along the component 2

**Supplementary Fig. 10:**
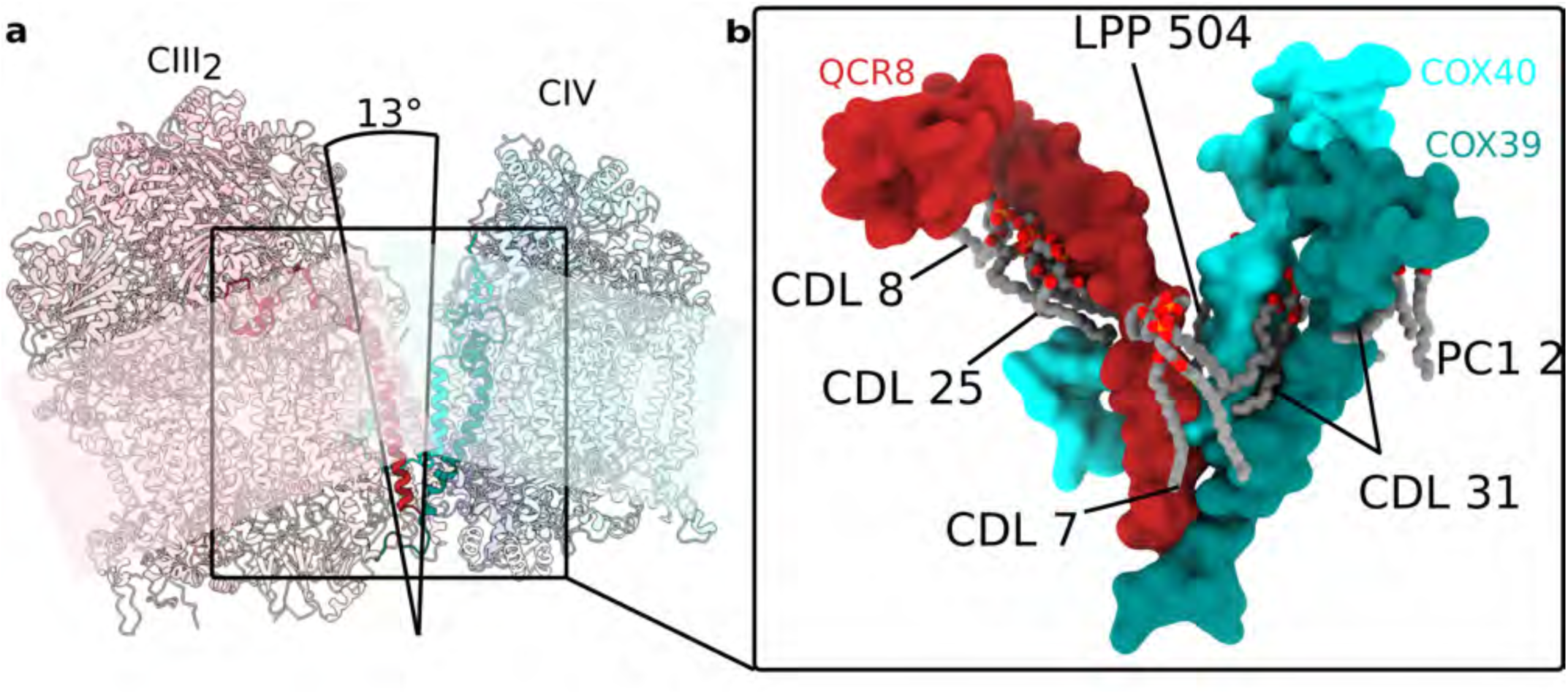
Lipids interaction with COX39, COX40, and QCR8. **a.** The position of COX39, COX40, and QCR8 with lipids in the consensus structure, in which 13° angle formed by the V-shape association of CIII_2_ and CIV in the consensus map. The membrane planes are indicated in the background. **b.** Lipids tightly interact with QCR8, COX39, and COX40 are shown.

**Supplementary Fig. 11:**
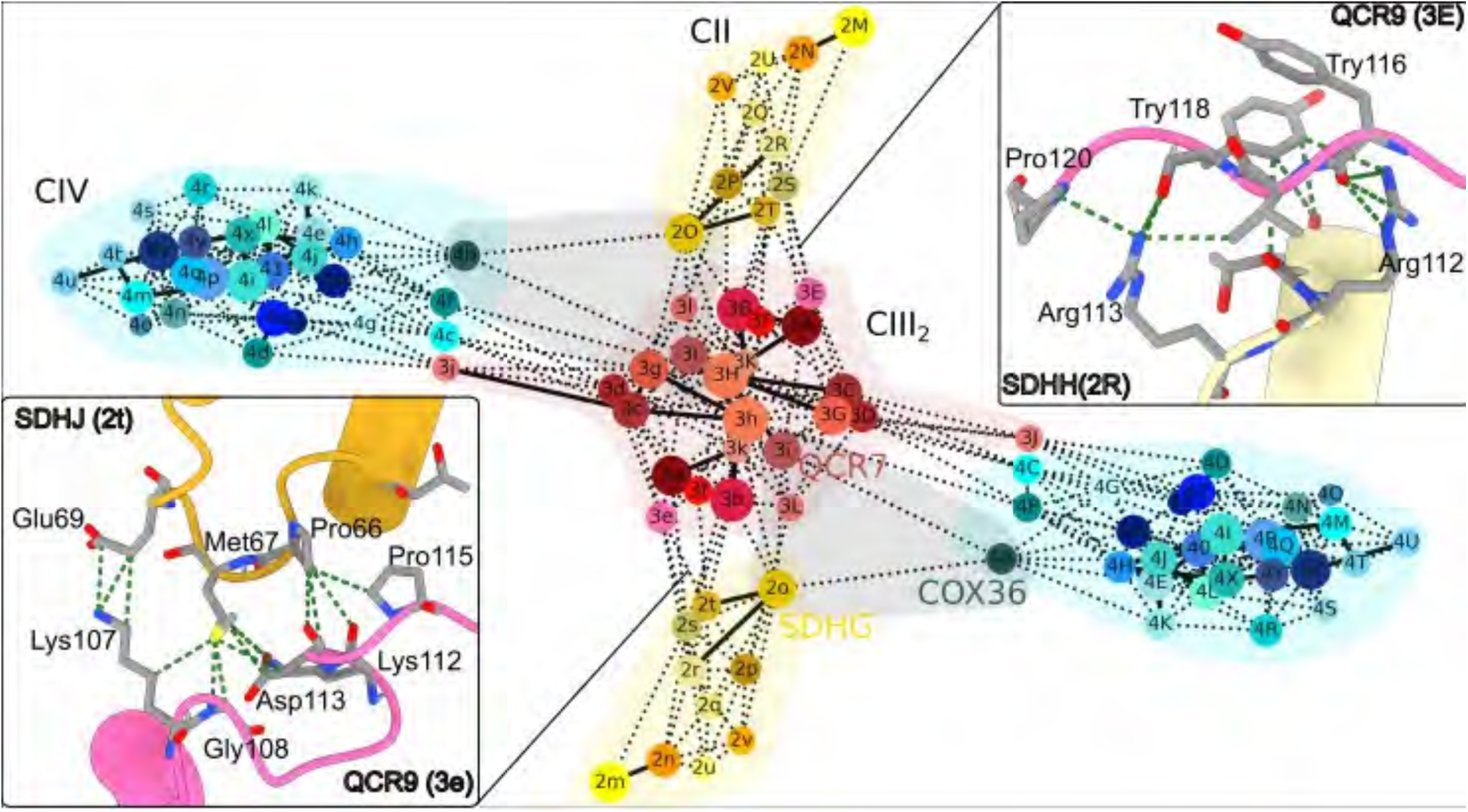
Protein interaction map between complexes. Schematic of protein-protein interactions, where nodes correspond to individual proteins (chain IDs indicated) and colored in shades of yellow (CII), red (CIII), cyan (CIV). The interactions were calculated using probe radius 3 Å using ChimeraX (Goddard *et al.* 2018).

**Supplementary Fig. 12:**
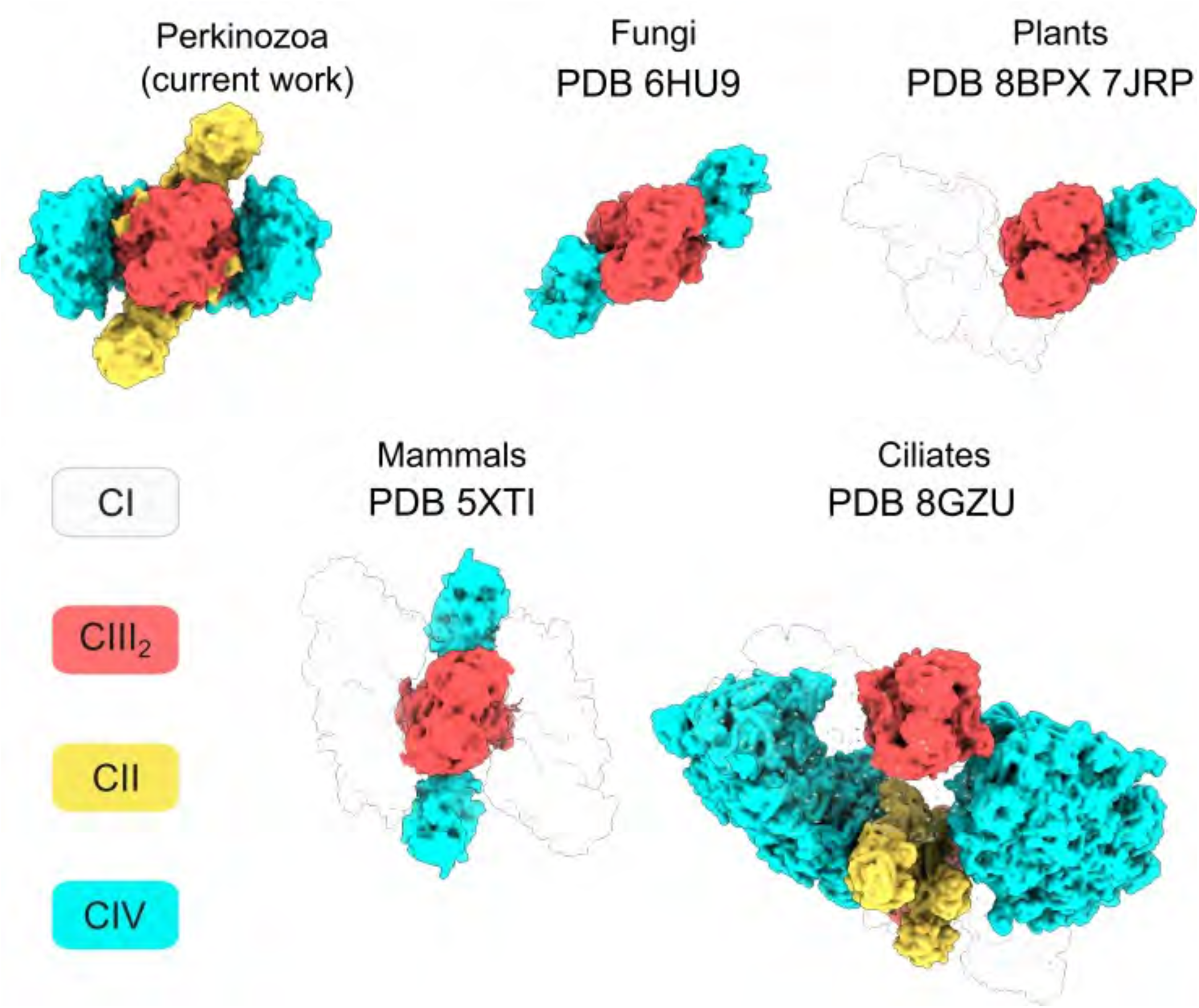
Diversity of respiratory SCs. Comparison of known modes between SCs from Perkinsozoa, fungi, plants, mammals, and ciliates. All contain CIII_2_ (red) and are shown in the same CIII orientation. PDB IDs of the models used are indicated.

**Supplementary Fig. 13:**
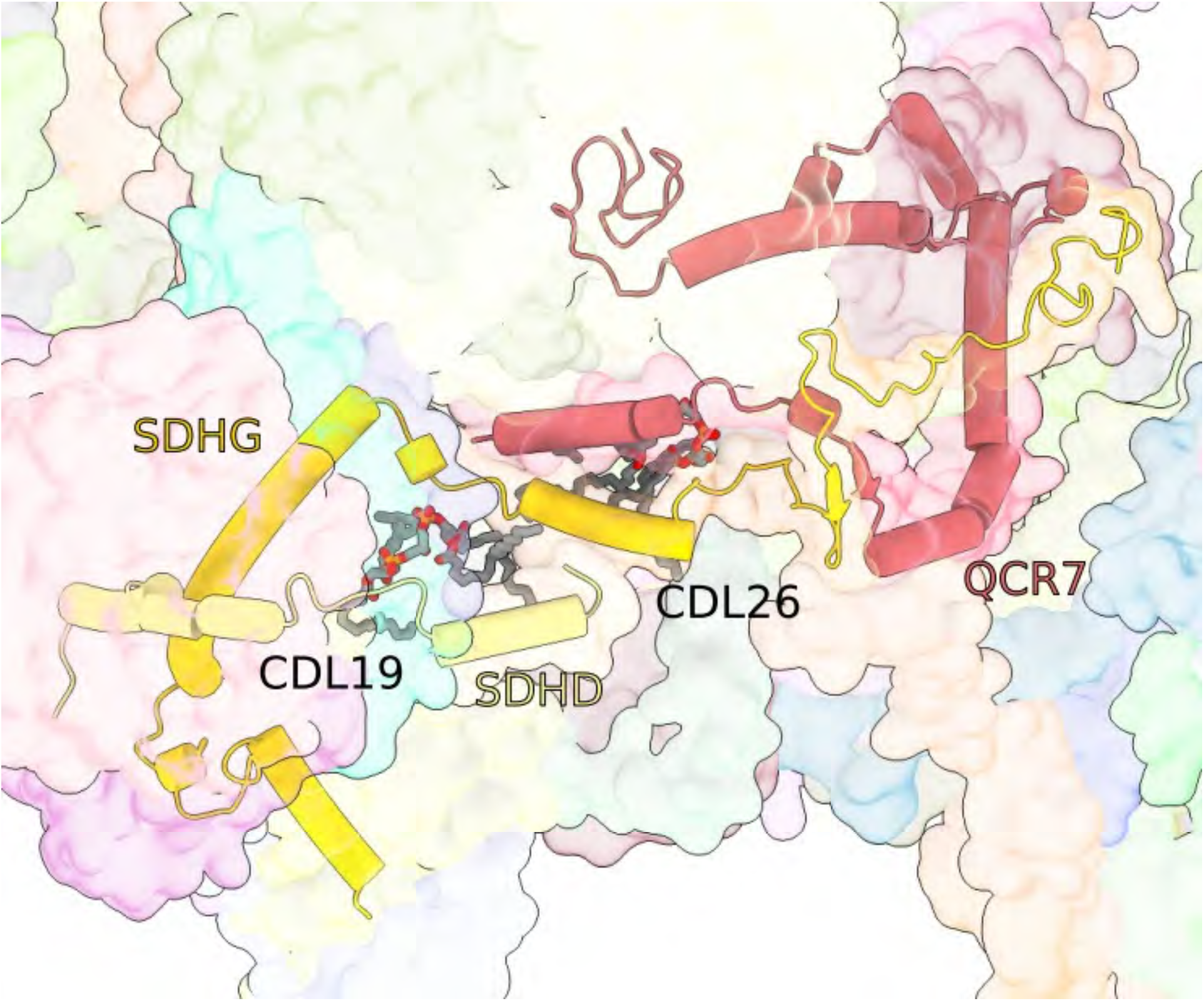
Protein lipids interactions at the II:III interface. Lipids CDL19 and CDL26 interact with SDHD and SHDG of CII and QCR7 of CIII, highlighting cardiolipin impliation in the SC formation.

**Supplementary Fig. 14:**
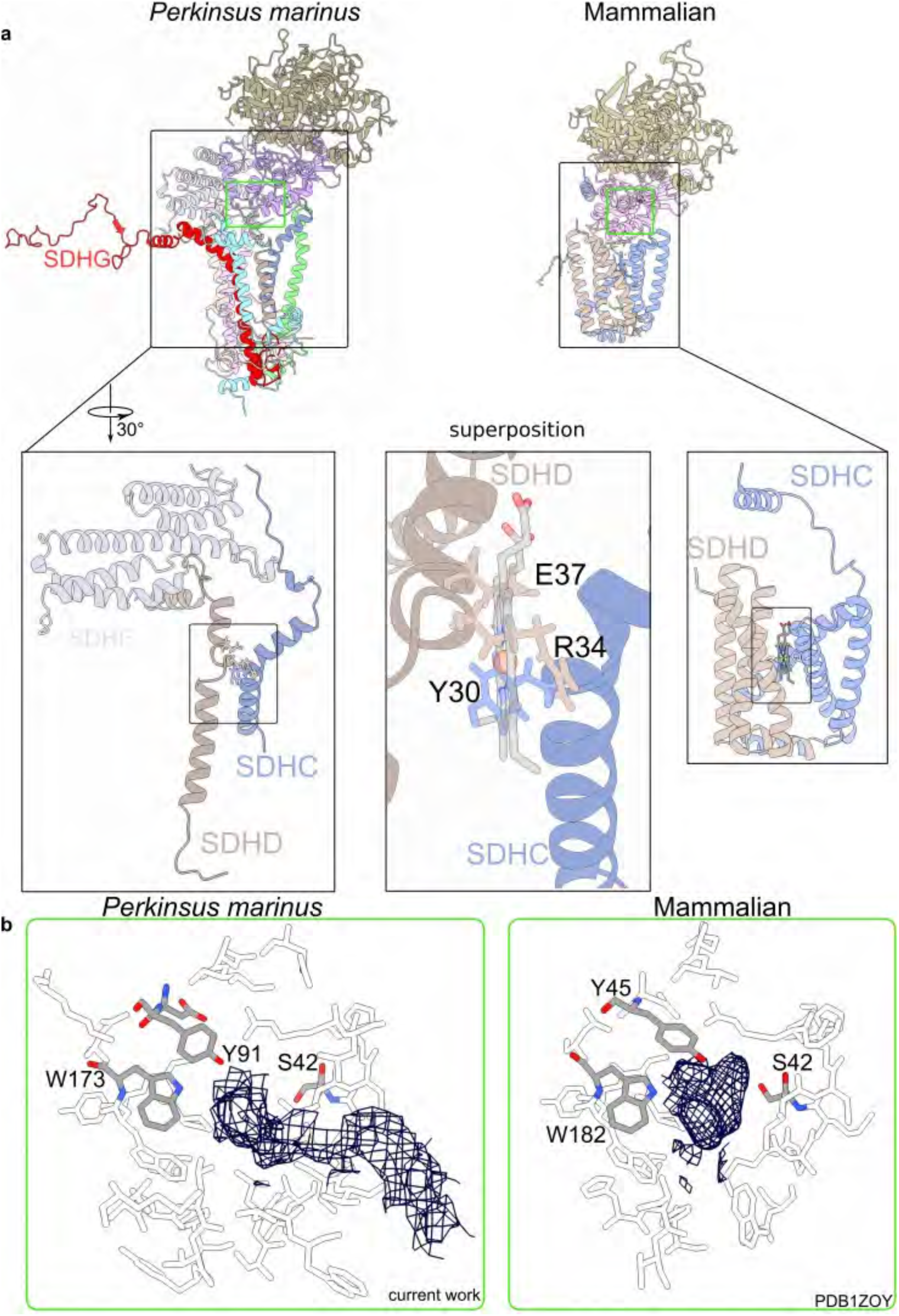
Subunits and interactions in CII. **a.** Comparison of CII structures between *Perkinsus* (current work) and mammals (PDB 1ZOY). Close-up views show the membrane-anchoring module SDHC-SDHD. SDHC-SDHD had not been not previously identified in Apicomplexa due to the divergent and truncated sequences. The superposition shows the the helices that contribute axial ligands to the heme group in mammals adopt a different conformation in *Perkinsus*, and that the prosthetic group is not compatible with our structure. **b.** Conserved residues in the quinone binding site of CII. Density for quinone is represented in black mesh in the site.

**Supplementary Fig. 15:**
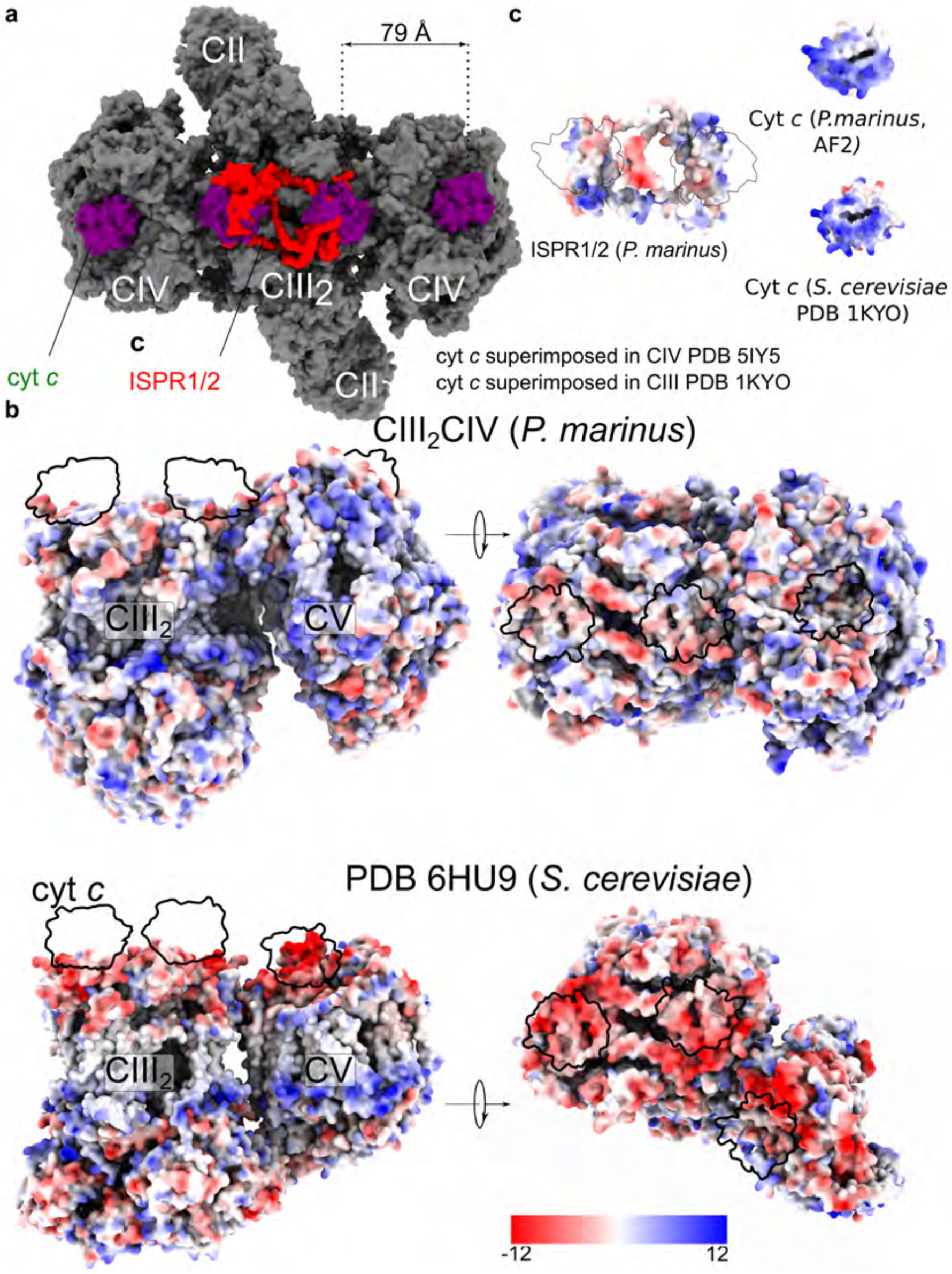
Electrostatic surface potential of SC, ISPR1,2 and Cyt *c*. **a.** Clashes between Cyt *c* and ISPR1,2 on the surface of the SC. The distance of the docking site for Cyt *c* between CIII and CIV is 79 Å. **b.** Comparison of the relative static electron potential surface of CIII-CIV between *P. marinus* and *S. cerevisiae*. **c.** Comparison of the surface of ISPR1,2 and Cyt *c* that interacts with CIII.

**Supplementary Fig. 16:**
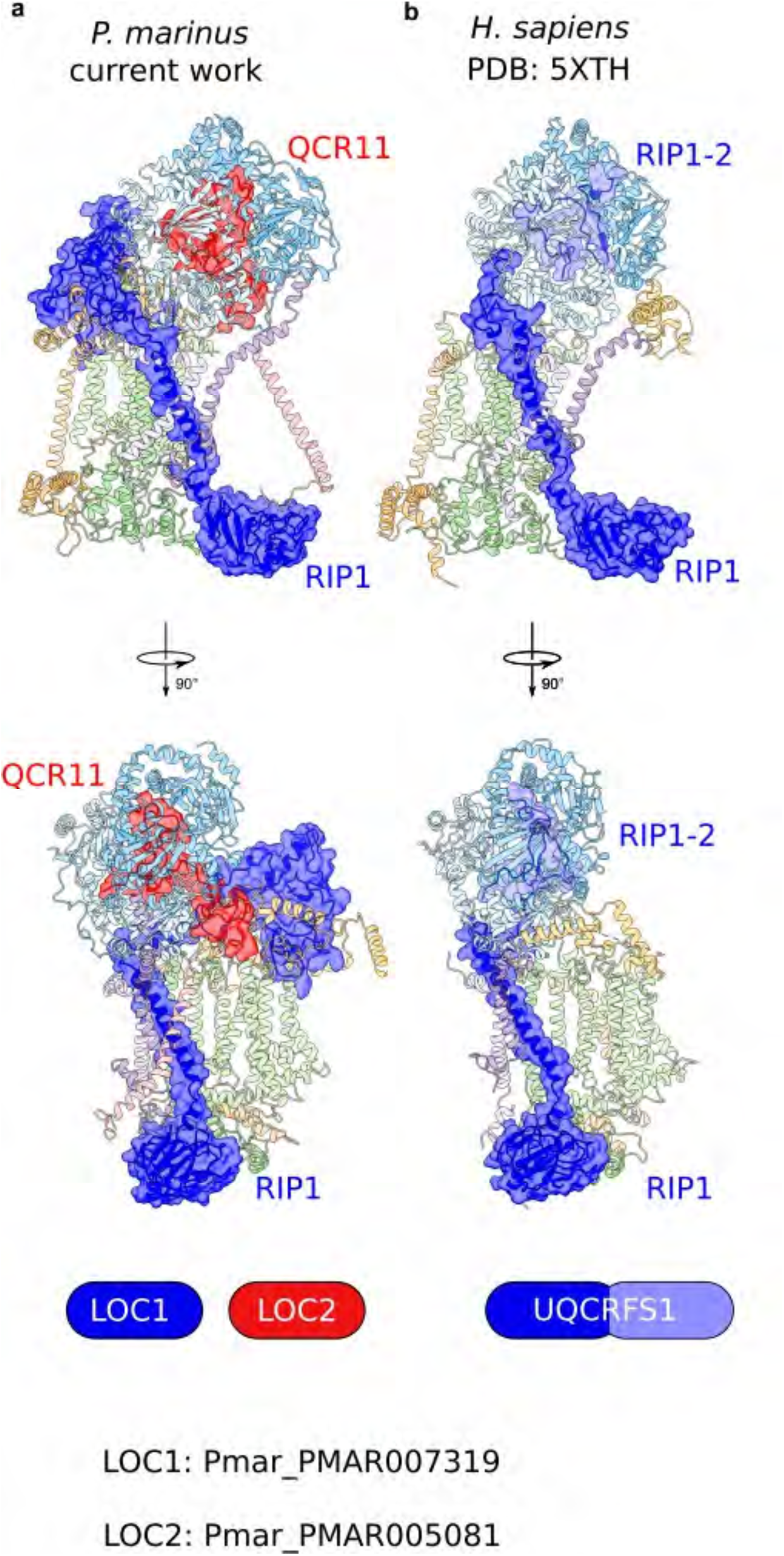
Parasite-specific subunit QCR11 substitutes the mammalian N-terminal partner RIP1-2. **a.** *P. marinus* RIP1 is blue, QCR11 is red. **b.** *H. Sapiens* RIP1 is blue, RIP1-2 is light blue.

**Supplementary Fig. 17:**
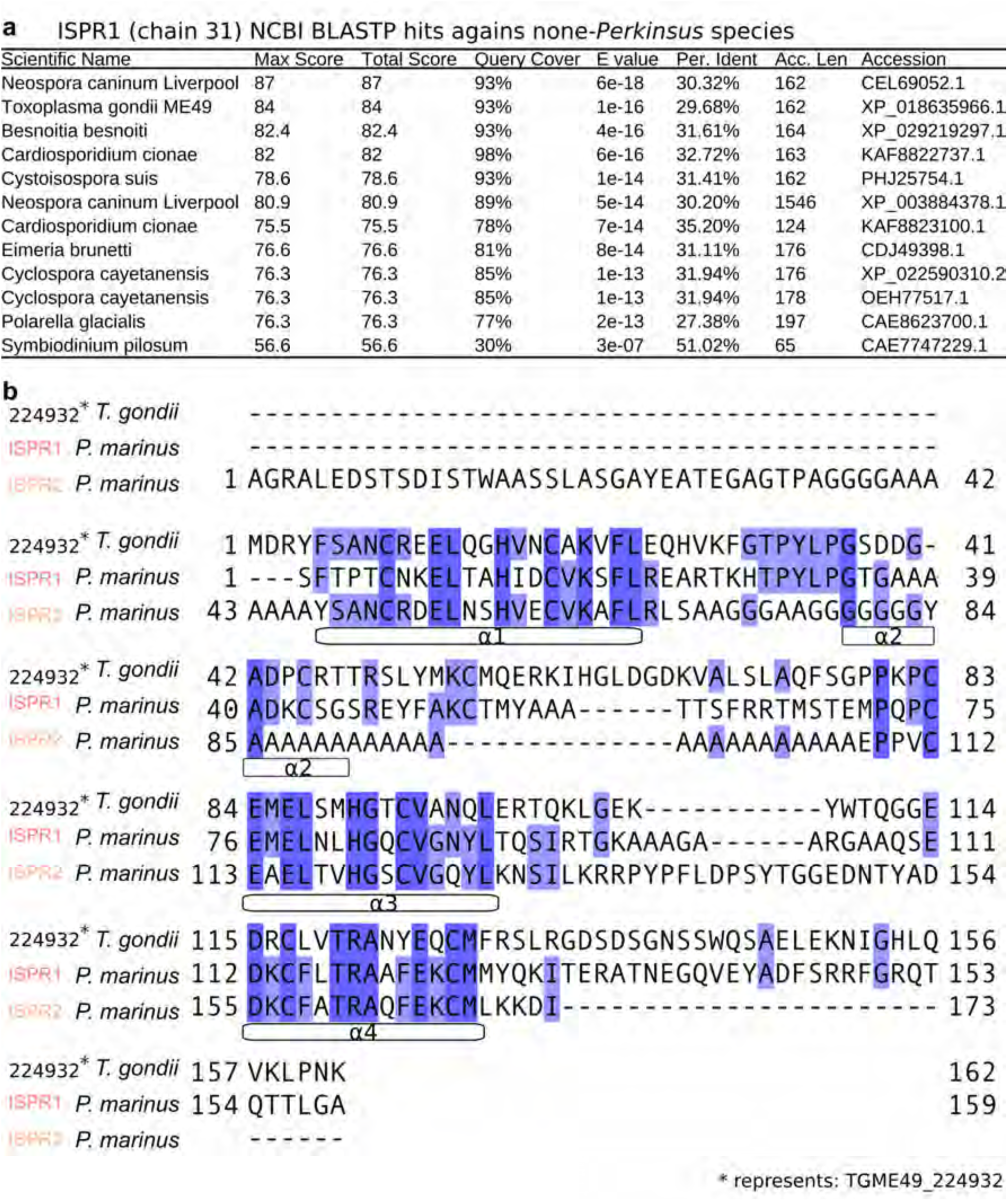
BLASTP hits for ISPR1 and identification of a putative ISPR2 homologue in *T. gondii*. **a.** Homologs of ISPR1 and ISPR2 in *Symbiodinium*, *Cyclospora*, *Toxoplasma* and *Neospora* species. **b.** The sequence alignment of *T. gondii* A0A125YMP1 with ISPR1, ISPR2, and. A0A125YMP1 and ISPR2 have C-terminal extensions.

**Supplementary Fig. 18:**
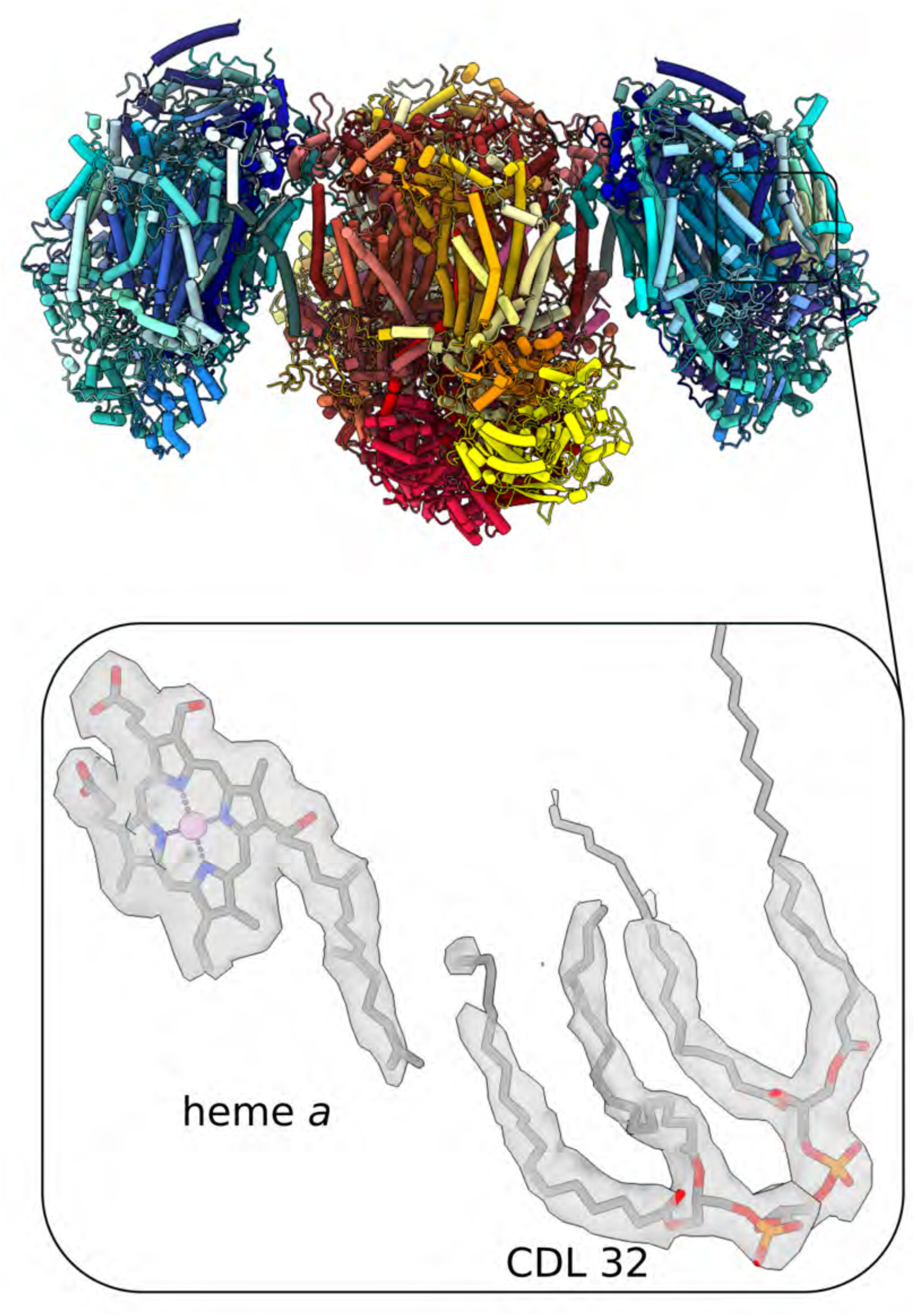
Interactions between the farnesyl tail of heme *a* and the tail of CDL 32 in CIV. The long hydrocarbon chain of heme a helps anchoring heme a into the hydrophobic environment.

**Supplementary Fig. 19:**
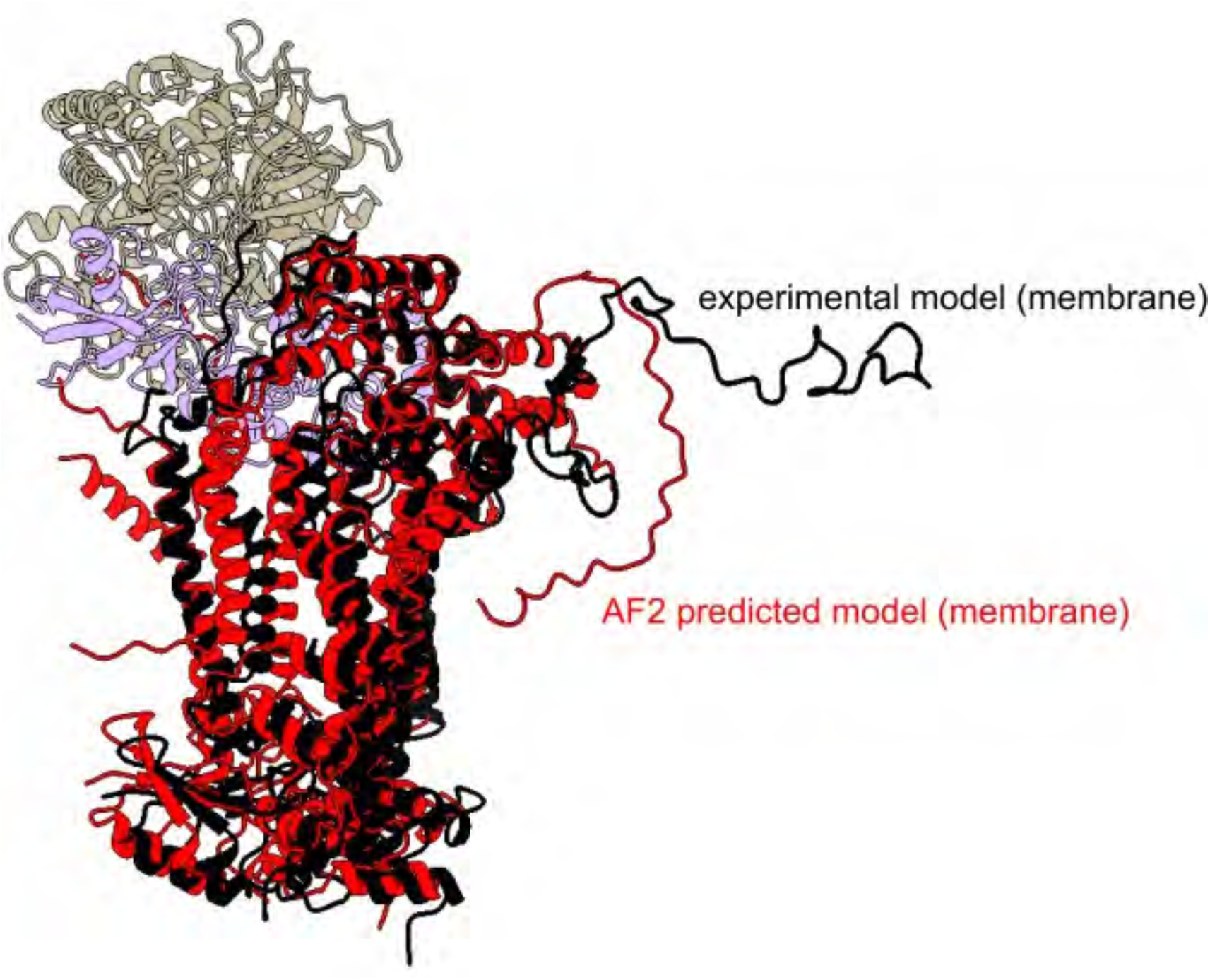
*AlphaFold2*-predicted transmembrane domain of CII. The cryo-EM determined structure is shown in black cartoon and the predicted model in red.

**Supplementary Fig. 20:**
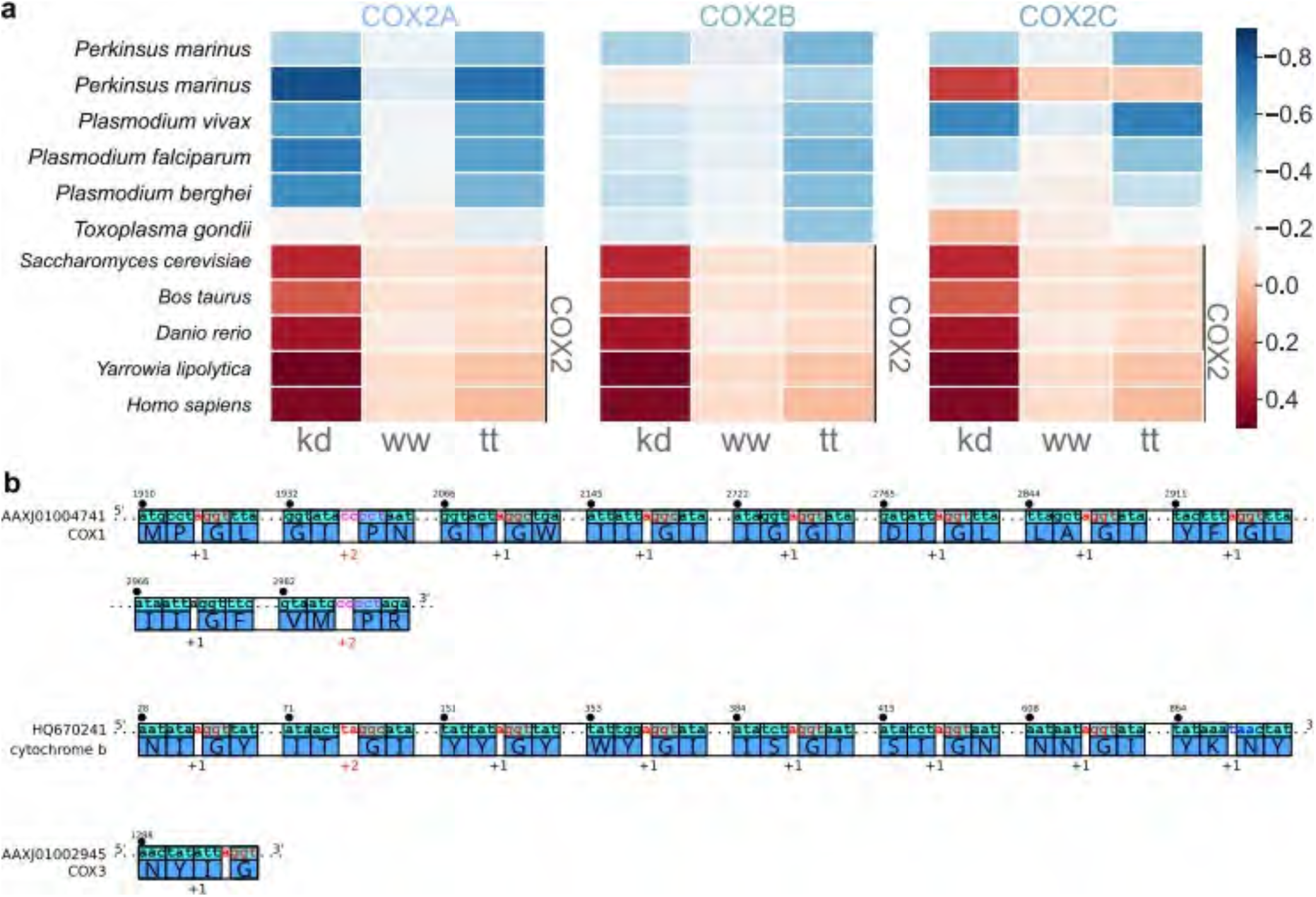
Protein splitting in COX2 and frameshifts in the mitochondria-encoded proteins COX1, Cyt *b*, COX3. **a.** Heat map indicating the average hydrophobicity of COX2 in divergent organisms, calculated as the grand average of hydropathy according to the Kyte-J.&Doolittle (kd) (Kyte and Doolittle, 1982), Moon-Fleming (mf) (Moon and Fleming, 2011) or Wimley-White (ww) (Wimley and White, 1996). **b.** Frameshifts found in mitochondria encoding three proteins COX1, COX3, and Cyt *b* with corresponding nucleotide and protein sequences.

**Table S1:**
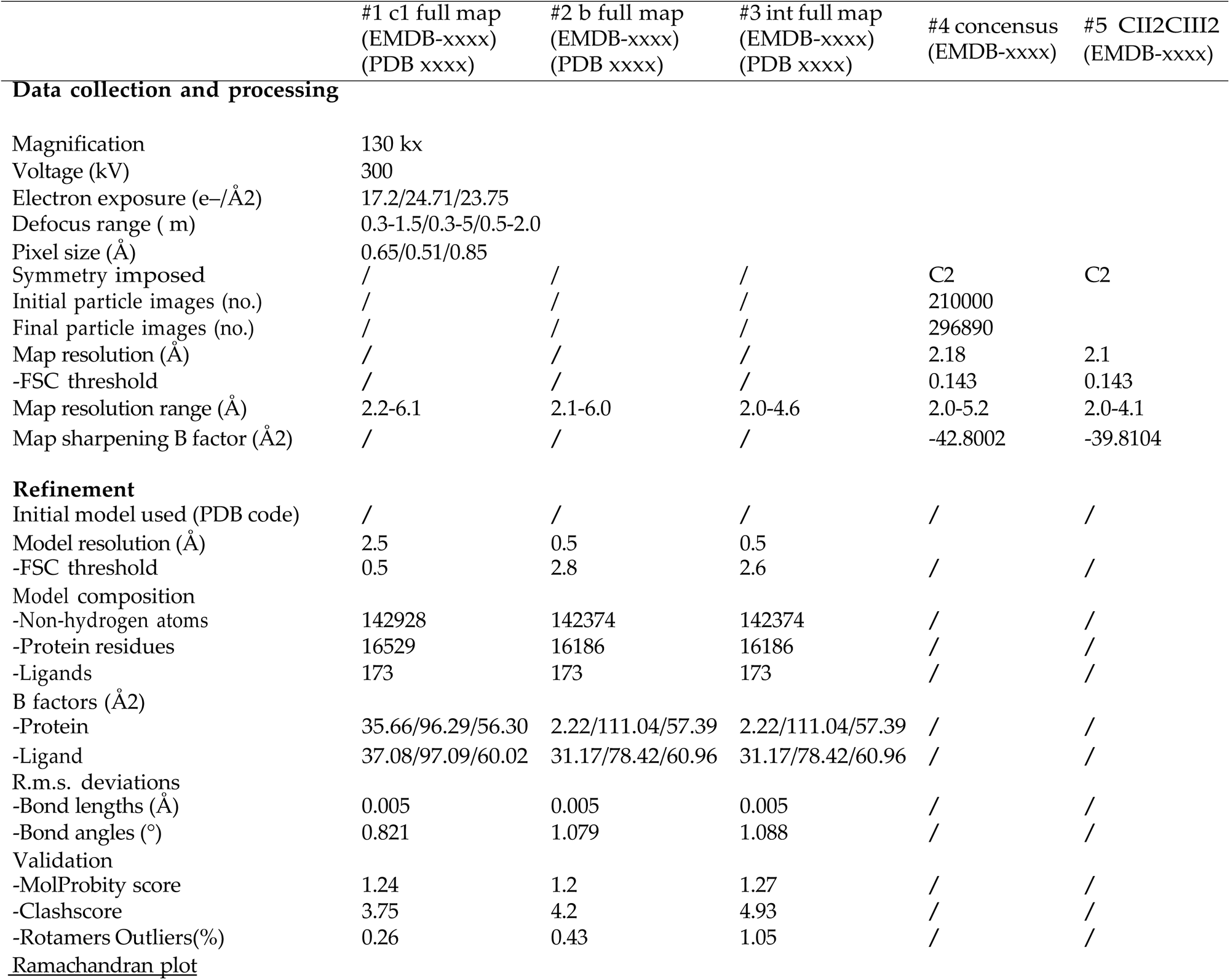
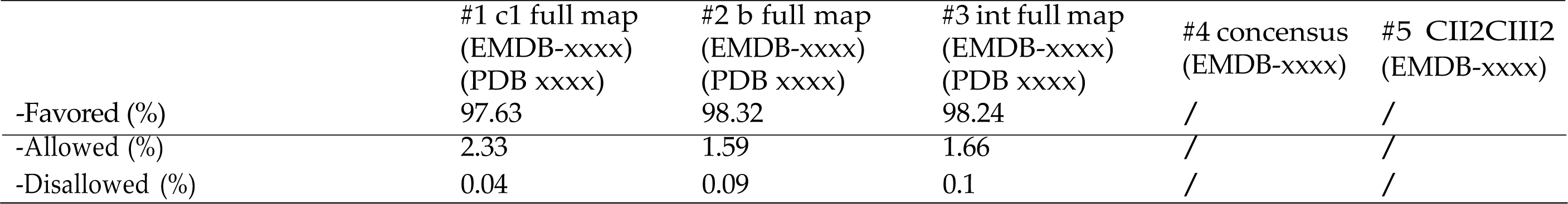
Cryo-EM data collection, refinement, and validation statistics (part 1)

**Table S2:** Cryo-EM data collection, refinement, and validation statistics (part 2)

|  | #6 CIV<br>(EMDB-xxxx) | #7 CII head<br>(EMDB-xxxx) | #8 c1 CII2CIII2<br>(EMDB-xxxx) | #9 b CII2CIII2<br>(EMDB-xxxx) | #10 int CII2CIII2<br>(EMDB-xxxx) |
| --- | --- | --- | --- | --- | --- |
| **Data collection and processing** |  |  |  |  |  |
| Magnification | 130 kx |  |  |  |  |
| Voltage (kV) | 300 |  |  |  |  |
| Electron exposure (e-/Å<sup>2</sup>) | 17.2/24.71/23.75 |  |  |  |  |
| Defocus range ( m) | 0.3-1.5/0.3-5/0.5-2.0 |  |  |  |  |
| Pixel size (Å) | 0.65/0.51/0.85 |  |  |  |  |
| Symmetry imposed | C1 | C1 | C1 | C2 | C2 |
| Initial particle images (no.) | 210000 |  |  |  |  |
| Final particle images (no.) | 296890 |  | 111717 | 47561 |  |
| Map resolution (Å) | 2.2 | 2.2 | 2.5 | 2.5 | 2.2 |
| FSC threshold | 0.143 | 0.143 | 0.143 | 0.143 | 0.143 |
| Map resolution range (Å) | 2.1-4.7 | 2.1-3.2 | 2.2-6.1 | 2.1-6.0 | 2.0-4.6 |
| Map sharpening B factor (Å<sup>2</sup>) | -32.6517 | -51.3825 | -31.9754 | -29.3593 | -31.7066 |

**Table S3:**
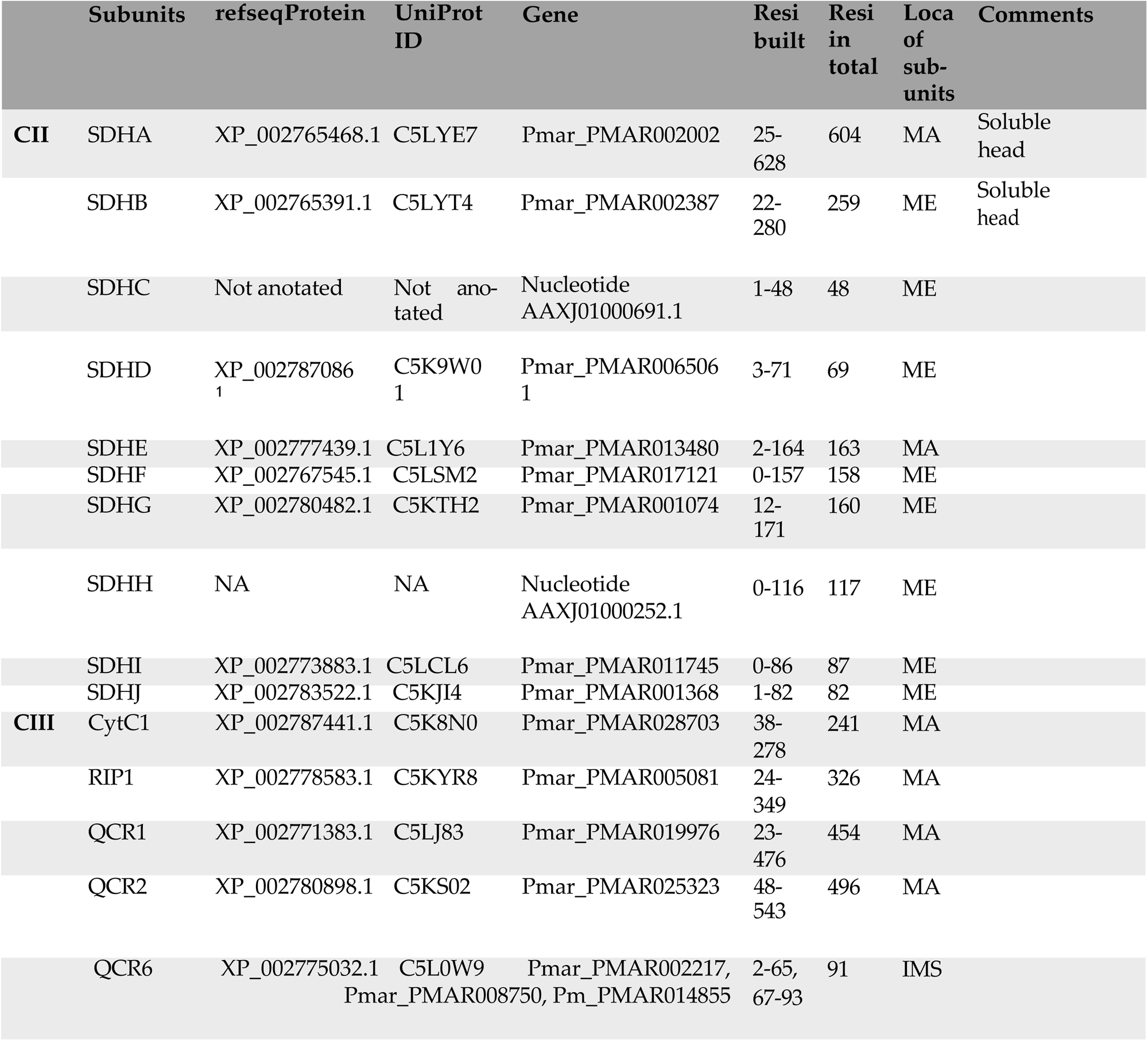
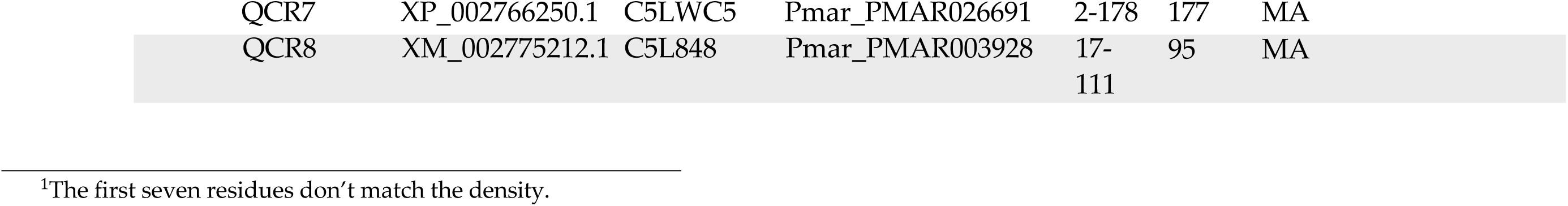
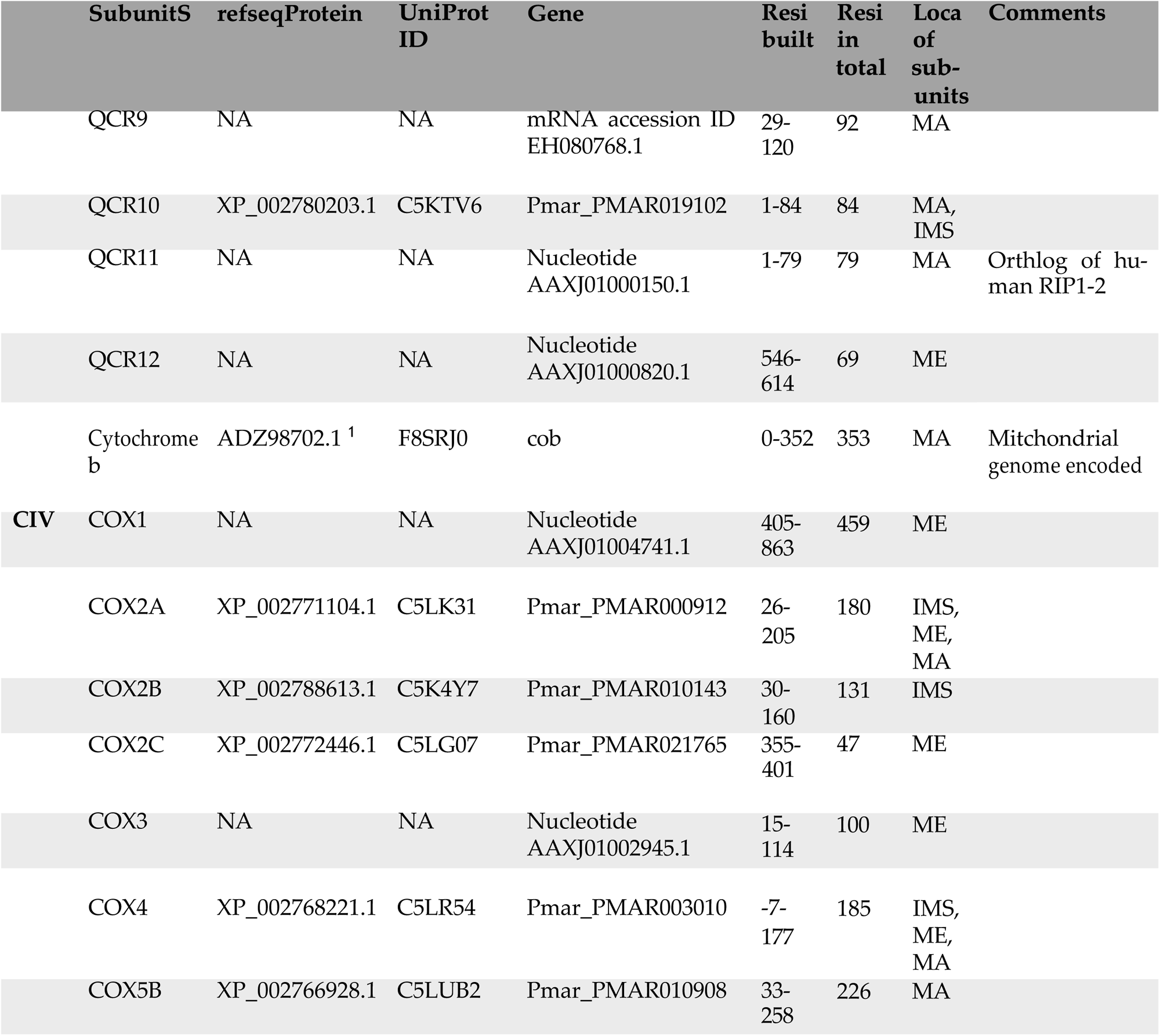
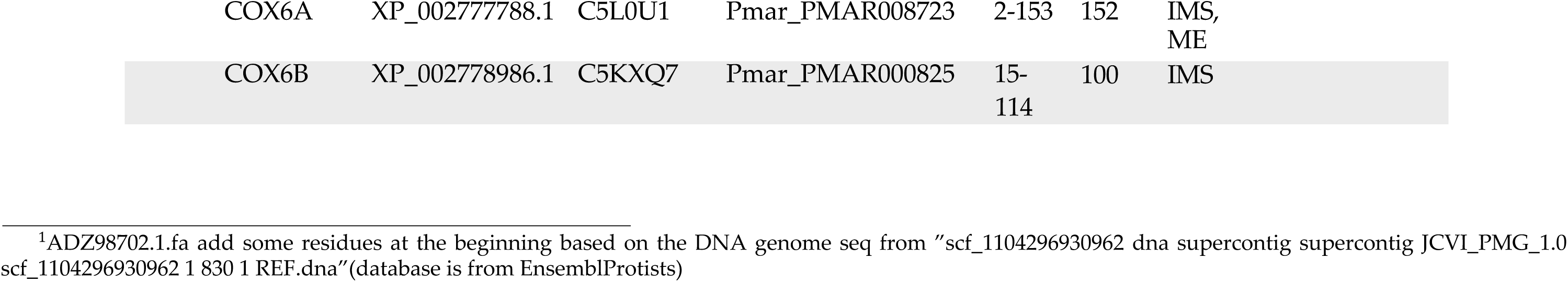
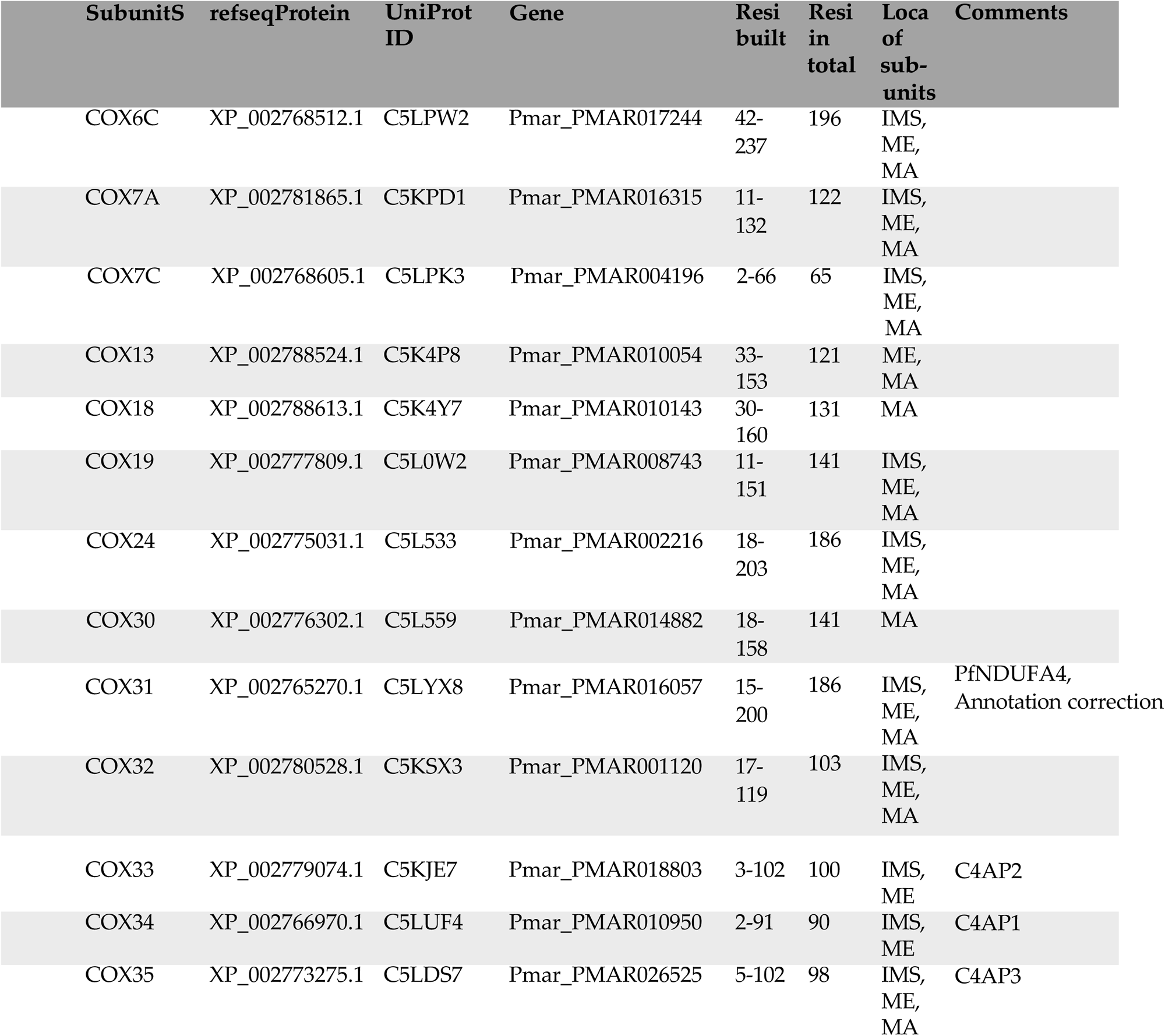
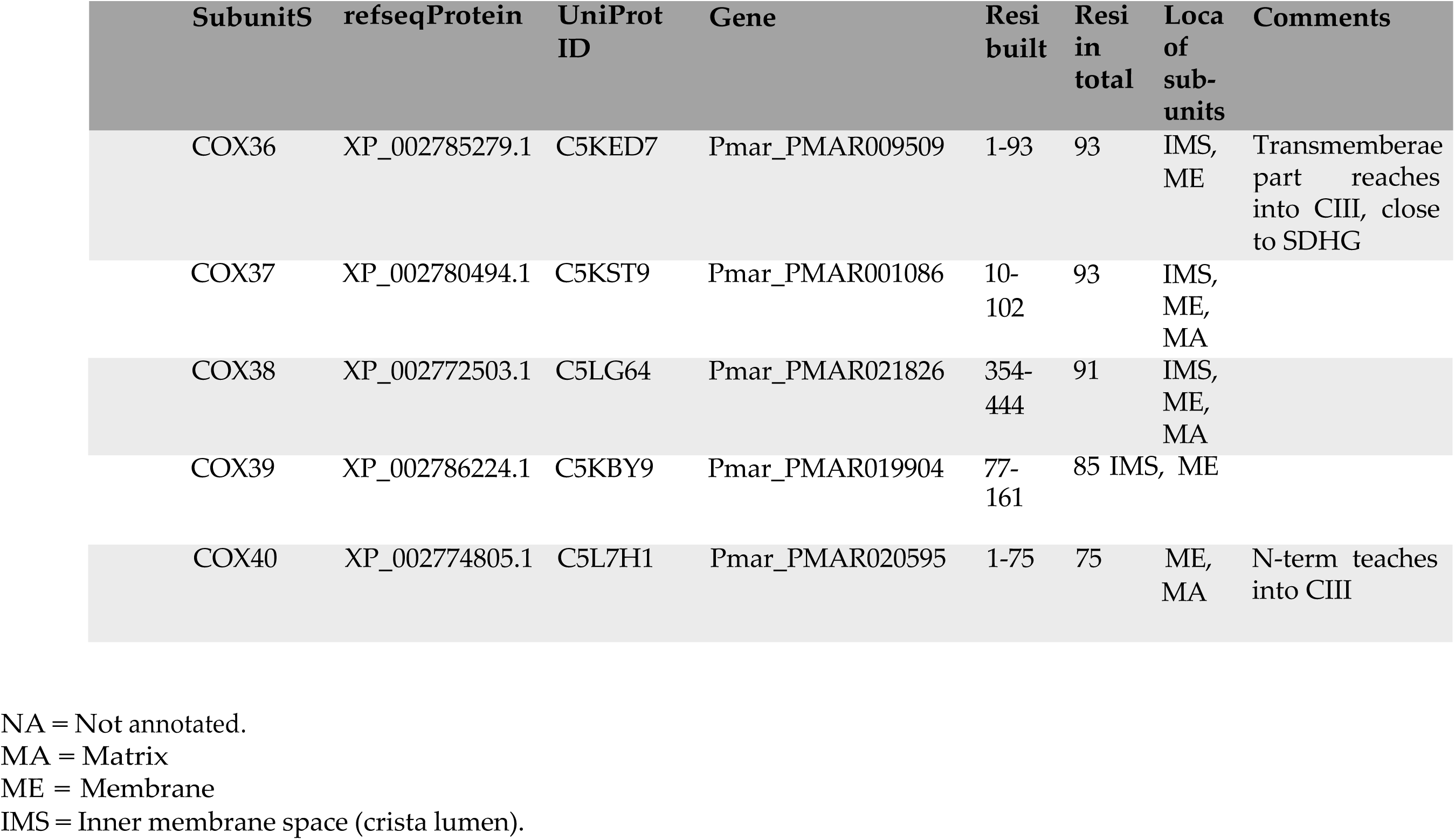
Model building statistics by subunits.

|  | Subunits | refseqProtein | UniProt ID | Gene | Resi built | Resi in total | Loca of sub-units | Comments |
| --- | --- | --- | --- | --- | --- | --- | --- | --- |
| **CII** | SDHA | XP_002765468.1 | C5LYE7 | Pmar_PMAR002002 | 25-628 | 604 | MA | Soluble head |
|  | SDHB | XP_002765391.1 | C5LYT4 | Pmar_PMAR002387 | 22-280 | 259 | ME | Soluble head |
|  | SDHC | Not anotated | Not anotated | Nucleotide AAXJ01000691.1 | 1-48 | 48 | ME |  |
|  | SDHD | XP_002787086<br>1 | C5K9W0<br>1 | Pmar_PMAR006506<br>1 | 3-71 | 69 | ME |  |
|  | SDHE | XP_002777439.1 | C5L1Y6 | Pmar_PMAR013480 | 2-164 | 163 | MA |  |
|  | SDHF | XP_002767545.1 | C5LSM2 | Pmar_PMAR017121 | 0-157 | 158 | ME |  |
|  | SDHG | XP_002780482.1 | C5KTH2 | Pmar_PMAR001074 | 12-171 | 160 | ME |  |
|  | SDHH | NA | NA | Nucleotide AAXJ01000252.1 | 0-116 | 117 | ME |  |
|  | SDHI | XP_002773883.1 | C5LCL6 | Pmar_PMAR011745 | 0-86 | 87 | ME |  |
|  | SDHJ | XP_002783522.1 | C5KJI4 | Pmar_PMAR001368 | 1-82 | 82 | ME |  |
| **CIII** | CytC1 | XP_002787441.1 | C5K8N0 | Pmar_PMAR028703 | 38-278 | 241 | MA |  |
|  | RIP1 | XP_002778583.1 | C5KYR8 | Pmar_PMAR005081 | 24-349 | 326 | MA |  |
|  | QCR1 | XP_002771383.1 | C5LJ83 | Pmar_PMAR019976 | 23-476 | 454 | MA |  |
|  | QCR2 | XP_002780898.1 | C5KS02 | Pmar_PMAR025323 | 48-543 | 496 | MA |  |
|  | QCR6 | XP_002775032.1 | C5L0W9 | Pmar_PMAR002217,<br>Pmar_PMAR008750, Pm_PMAR014855 | 2-65,<br>67-93 | 91 | IMS |  |

|  |  |  |  |  |  |  |
| --- | --- | --- | --- | --- | --- | --- |
| QCR7 | XP_002766250.1 | C5LWC5 | Pmar_PMAR026691 | 2-178 | 177 | MA |
| QCR8 | XM_002775212.1 | C5L848 | Pmar_PMAR003928 | 17-111 | 95 | MA |

|  | SubunitS | refseqProtein | UniProt ID | Gene | Resi built | Resi in total | Loca of sub-units | Comments |
| --- | --- | --- | --- | --- | --- | --- | --- | --- |
|  | QCR9 | NA | NA | mRNA accession ID EH080768.1 | 29-120 | 92 | MA |  |
|  | QCR10 | XP_002780203.1 | C5KTV6 | Pmar_PMAR019102 | 1-84 | 84 | MA, IMS |  |
|  | QCR11 | NA | NA | Nucleotide AAXJ01000150.1 | 1-79 | 79 | MA | Orthlog of human RIP1-2 |
|  | QCR12 | NA | NA | Nucleotide AAXJ01000820.1 | 546-614 | 69 | ME |  |
|  | Cytochrome b | ADZ98702.1 <sup>1</sup> | F8SRJ0 | cob | 0-352 | 353 | MA | Mitochondrial genome encoded |
| **CIV** | COX1 | NA | NA | Nucleotide AAXJ01004741.1 | 405-863 | 459 | ME |  |
|  | COX2A | XP_002771104.1 | C5LK31 | Pmar_PMAR000912 | 26-205 | 180 | IMS, ME, MA |  |
|  | COX2B | XP_002788613.1 | C5K4Y7 | Pmar_PMAR010143 | 30-160 | 131 | IMS |  |
|  | COX2C | XP_002772446.1 | C5LG07 | Pmar_PMAR021765 | 355-401 | 47 | ME |  |
|  | COX3 | NA | NA | Nucleotide AAXJ01002945.1 | 15-114 | 100 | ME |  |
|  | COX4 | XP_002768221.1 | C5LR54 | Pmar_PMAR003010 | -7-177 | 185 | IMS, ME, MA |  |
|  | COX5B | XP_002766928.1 | C5LUB2 | Pmar_PMAR010908 | 33-258 | 226 | MA |  |

|  |  |  |  |  |  |  |
| --- | --- | --- | --- | --- | --- | --- |
| COX6A | XP_002777788.1 | C5L0U1 | Pmar_PMAR008723 | 2-153 | 152 | IMS,<br>ME |
| COX6B | XP_002778986.1 | C5KXQ7 | Pmar_PMAR000825 | 15-114 | 100 | IMS |

| SubunitS | refseqProtein | UniProt ID | Gene | Resi built | Resi in total | Loca of sub-units | Comments |
| --- | --- | --- | --- | --- | --- | --- | --- |
| COX6C | XP_002768512.1 | C5LPW2 | Pmar_PMAR017244 | 42-237 | 196 | IMS, ME, MA |  |
| COX7A | XP_002781865.1 | C5KPD1 | Pmar_PMAR016315 | 11-132 | 122 | IMS, ME, MA |  |
| COX7C | XP_002768605.1 | C5LPK3 | Pmar_PMAR004196 | 2-66 | 65 | IMS, ME, MA |  |
| COX13 | XP_002788524.1 | C5K4P8 | Pmar_PMAR010054 | 33-153 | 121 | ME, MA |  |
| COX18 | XP_002788613.1 | C5K4Y7 | Pmar_PMAR010143 | 30-160 | 131 | MA |  |
| COX19 | XP_002777809.1 | C5L0W2 | Pmar_PMAR008743 | 11-151 | 141 | IMS, ME, MA |  |
| COX24 | XP_002775031.1 | C5L533 | Pmar_PMAR002216 | 18-203 | 186 | IMS, ME, MA |  |
| COX30 | XP_002776302.1 | C5L559 | Pmar_PMAR014882 | 18-158 | 141 | MA |  |
| COX31 | XP_002765270.1 | C5LYX8 | Pmar_PMAR016057 | 15-200 | 186 | IMS, ME, MA | PfNDUFA4, Annotation correction |
| COX32 | XP_002780528.1 | C5KSX3 | Pmar_PMAR001120 | 17-119 | 103 | IMS, ME, MA |  |
| COX33 | XP_002779074.1 | C5KJE7 | Pmar_PMAR018803 | 3-102 | 100 | IMS, ME | C4AP2 |
| COX34 | XP_002766970.1 | C5LUF4 | Pmar_PMAR010950 | 2-91 | 90 | IMS, ME | C4AP1 |
| COX35 | XP_002773275.1 | C5LDS7 | Pmar_PMAR026525 | 5-102 | 98 | IMS, ME, MA | C4AP3 |
| COX36 | XP_002785279.1 | C5KED7 | Pmar_PMAR009509 | 1-93 | 93 | IMS, ME | Transmembrerae part reaches into CIII, close to SDHG |
| COX37 | XP_002780494.1 | C5KST9 | Pmar_PMAR001086 | 10-102 | 93 | IMS, ME, MA |  |
| COX38 | XP_002772503.1 | C5LG64 | Pmar_PMAR021826 | 354-444 | 91 | IMS, ME, MA |  |
| COX39 | XP_002786224.1 | C5KBY9 | Pmar_PMAR019904 | 77-161 | 85 | IMS, ME |  |
| COX40 | XP_002774805.1 | C5L7H1 | Pmar_PMAR020595 | 1-75 | 75 | ME, MA | N-term teaches into CIII |

NA = Not annotated.
MA = Matrix
ME = Membrane
IMS = Inner membrane space (crista lumen).

